# Protracted yet coordinated differentiation of long-lived SARS-CoV-2-specific CD8+ T cells during COVID-19 convalescence

**DOI:** 10.1101/2021.04.28.441880

**Authors:** Tongcui Ma, Heeju Ryu, Matthew McGregor, Benjamin Babcock, Jason Neidleman, Guorui Xie, Ashley F. George, Julie Frouard, Victoria Murray, Gurjot Gill, Eliver Ghosn, Evan Newell, Sulggi Lee, Nadia R. Roan

**Affiliations:** Gladstone Institutes, San Francisco, CA, USA; Department of Urology, University of California, San Francisco, CA, USA; Vaccine and Infectious Disease Division Fred Hutchison Cancer Research Center, Seattle, WA, USA; Department of Medicine, Lowance Center for Human Immunology, Emory Vaccine Center, Emory University, Atlanta, GA, USA; Department of Pediatrics, Lowance Center for Human Immunology, Emory Vaccine Center, Emory University, Atlanta, GA, USA; Zuckerberg San Francisco General Hospital and the University of California, San Francisco, CA, USA

## Abstract

CD8+ T cells are important antiviral effectors that can potentiate long-lived immunity against COVID-19, but a detailed characterization of these cells has been hampered by technical challenges. We screened 21 well-characterized, longitudinally-sampled convalescent donors that recovered from mild COVID-19 against a collection of SARS-CoV-2 tetramers, and identified one participant with an immunodominant response against Nuc_322-331_, a peptide that is conserved in all the SARS-CoV-2 variants-of-concern reported to date. We conducted 38- parameter CyTOF phenotyping on tetramer-identified Nuc_322-331_-specific CD8+ T cells, and on CD4+ and CD8+ T cells recognizing the entire nucleocapsid and spike proteins from SARS-CoV-2, and took 32 serological measurements on longitudinal specimens from this participant. We discovered a coordination of the Nuc_322-331_-specific CD8+ T response with both the CD4+ T cell and antibody pillars of adaptive immunity. Nuc_322-331_-specific CD8+ T cells were predominantly central memory T cells, but continually evolved over a ∼6-month period of convalescence. We observed a slow and progressive decrease in the activation state and polyfunctionality of the Nuc_322-331_-specific CD8+ T cells, accompanied by an increase in their lymph-node homing and homeostatic proliferation potential. These results suggest that following a typical case of mild COVID-19, SARS-CoV-2-specific CD8+ T cells not only persist but continuously differentiate in a coordinated fashion well into convalescence, into a state characteristic of long-lived, self-renewing memory.

## INTRODUCTION

The uncertainty about the longevity of the immune response elicited by prior SARS-CoV- 2 infection or vaccination has been a major area of concern as the world tries to exit from the ongoing COVID-19 pandemic. Studies at the start of the pandemic suggesting a short-lived SARS-CoV-2 antibody response (1) brought about widespread concern, but follow-up studies now suggest that infected individuals exhibit a prolonged and evolving humoral immune response (2, 3). Furthermore, SARS-CoV-2-specific memory T cells – a second arm of adaptive immunity – can be detected more than six months into convalescence and these cells can self-renew in response to the homeostatic proliferation cytokine IL7 (4–6). Encouragingly, memory T cells against the nucleocapsid protein from the closely-related SARS-CoV-1 virus can be detected 17 years after infection (7), suggesting the potential for durable T cell immunity against pathogenic beta-coronaviruses. Importantly, relative to antibodies, T cells are less prone to evasion by the variants of concern emerging worldwide (8), suggesting a potentially important role for these immune effectors in long-term population-based immunity in the years ahead.

Characterizing the memory T cells responding to SARS-CoV-2 will improve our understanding of the features defining long-lived immunity, and of the ability of T cells to protect against reinfection. While the breadth of the SARS-CoV-2-specific response during convalescence has been extensively examined (9, 10), much less is known about the phenotypes of SARS-CoV-2-specific memory T cells. To phenotype SARS-CoV-2-specific T cells, most studies rely on stimulating T cells with SARS-CoV-2-specific antigens/peptides, and examining the cells that respond by expressing activation-induced markers (AIM) or cytokines (5, 9, 11–13). These studies likely underestimate the phenotypic complexity of antigen-specific T cells because of the limited number of AIMs or cytokine markers that can be used to identify responsive cells. These assays are also limited, because they do not capture antigen-specific T cells in their original, unstimulated states. Detecting antigen-specific unstimulated cells requires other, more technically-involved approaches, such as the use of T cell multimers/tetramers. Tetramers, which consist of four linked peptide-MHC complexes that specifically bind epitope-specific T cells, are one of the only ways to examine the original phenotypes of antigen-specific T cells. A handful of studies have incorporated the use of SARS-CoV-2 MHC class I multimers to examine CD8+ T cell responses (13–18). Due to small numbers of multimer+ cells of a single specificity, most of these studies examined the combined phenotypes of multimer+ cells recognizing different epitopes and/or pre-enriched for multimer+ cells (to increase detectability) which can bias the resulting collection of antigen-specific cells. One of the studies (17) conducted a longitudinal analysis multimer+ cells from one patient at 6 timepoints – 2 during acute infection and 4 at convalescence – by examining by FACS the levels of 5 phenotyping parameters on pre-enriched multimer+ cells. Although these studies together have revealed multimer+ cells to be distributed among multiple canonical subsets and pinpointed a handful of surface markers expressed by these cells, the inability to identify enough epitope-specific cells for high-parameter phenotypic analysis has made it challenging to perform a comprehensive analysis of how SARS-CoV-2- specific CD8+ T cells against a defined specificity evolve over the course of convalescence.

To fill this void, we screened banked longitudinal specimens from the UCSF COVID-19 Host Immune Response Pathogenesis (CHIRP) cohort against a collection of SARS-CoV-2 tetramers, to try to identify an immunodominant response that can be captured by tetramer analysis. This screen identified one individual with a particularly robust response detectable by one of the tetramers harboring a nucleocapsid peptide. The immunodominance of this response enabled us to perform a longitudinal analysis without the need to pre-enrich for tetramer+ cells or combine tetramer+ cells of multiple specificities. By combining 38-parameter CyTOF phenotyping with detection of these tetramer+ cells, we established an in-depth view of epitope-specific T cell responses at 5 longitudinal timepoints from early to late (>6 months) COVID-19 convalescence. Effector responses by these epitope-specific T cells were examined by treating cells with cognate peptide and examining by CyTOF the cytokine and cytolytic effector mechanisms of these cells. All longitudinal specimens were also phenotyped by the same effector CyTOF panel for total nucleocapsid- and spike-specific CD4+ and CD8+ T cell responses, and assessed for the levels of 32 different isotype-specific SARS-CoV-2 antibodies. All together, we measured nearly 400 different SARS-CoV-2-specific parameters for each of the 5 timepoints, and analyzed them in association with features of the tetramer+ response in order to provide an integrated and comprehensive overview of the immunological context surrounding the epitope-specific CD8+ T cell response. Although this study focuses on only one individual, this person exhibited a typical mild course of infection that was very well-defined clinically and that fully resolved in a timely manner. We therefore consider the response we have characterized to potentially reflect a common one in individuals that have recovered from mild COVID-19.

## RESULTS

### Identification and description of case study PID4103 with immunodominant Nuc_322-331_ CD8+ T cell response

The COVID-19 Host Immune Pathogenesis (CHIRP) study is a prospective longitudinal study designed to understand the evolution of host responses from the acute to convalescent phases of SARS-CoV-2 infection. Individuals within 31 days from symptom onset or SARS-CoV- 2 exposure were enrolled, and participants were sampled for 6 months. Cryopreserved PBMCs from a total of 21 convalescent CHIRP cohort participants from which specimens from 5 longitudinal study visits were available, and who had mild COVID-19 disease (see Methods), were screened by flow cytometry against 9 MHC class I tetramers harboring predicted CD8+ T cell epitopes from the spike and nucleocapsid proteins of SAR-CoV-2 (Table S1). We identified one donor, PID4103, who harbored an immunodominant response against an HLA-B*40:01-restricted peptide (sequence MEVTPSGTWL) spanning residues 322 to 311 of nucleocapsid (Nuc_322-331_) (Fig. 1A). This peptide is 100% conserved in the B.1.1.7, B.1.351, P.1, and B429/CAL20C variants of concern, as well as in the 2002 SARS-CoV-1 virus, but absent from the four common cold coronaviruses 229E, NL63, OC43, and HKU1 (Fig. S1).

**Figure 1.**
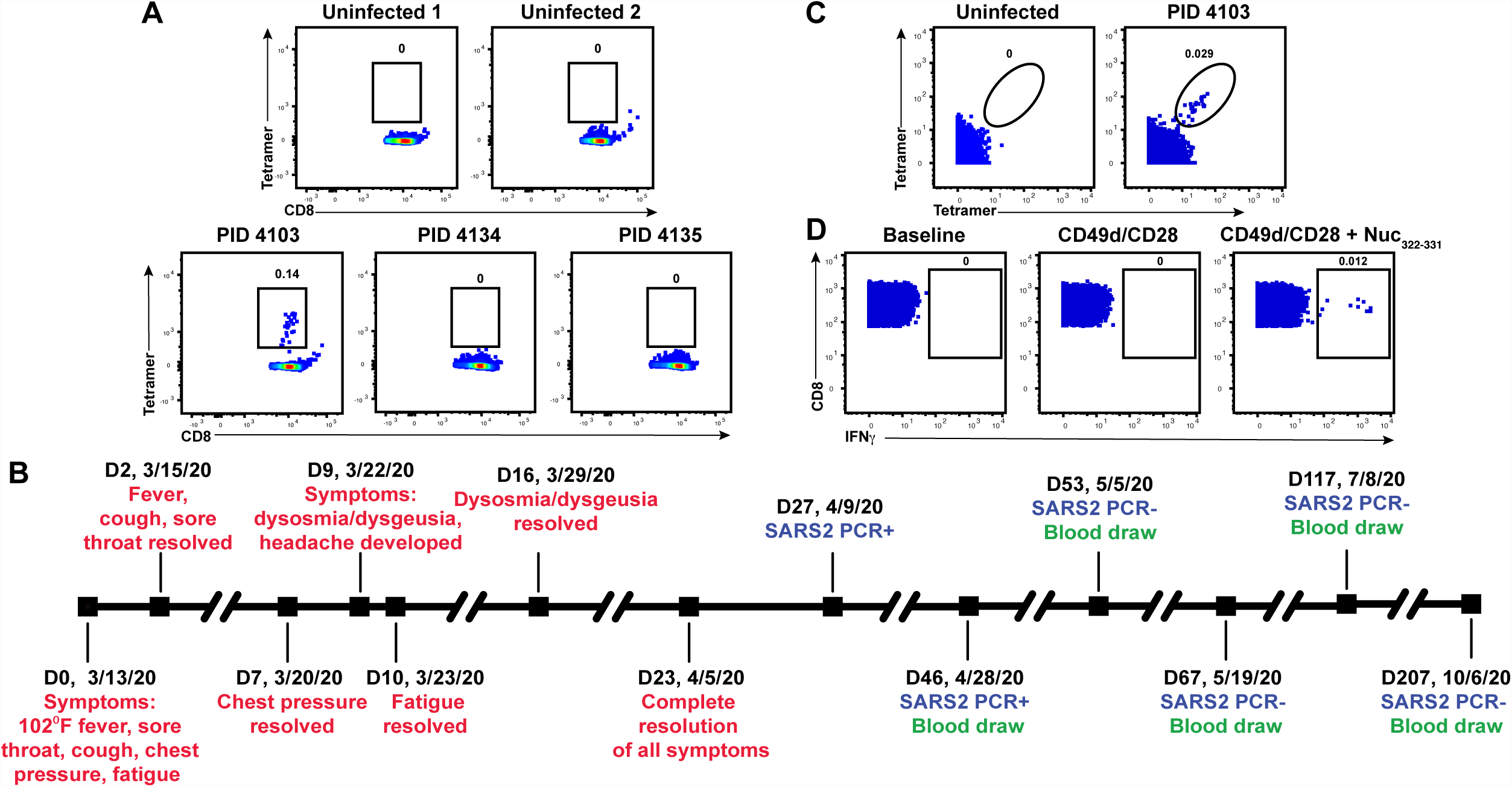
Identification and description of case study PID4103 with immunodominant CD8+ T cell response against Nuc_331-331_. **(A)** A distinct population of Nuc_322-331_-specific CD8+ T cells is detected by FACS tetramer staining in convalescent donor PID4103. *Top:* PBMCs from uninfected individuals were analyzed by FACS for binding to the HLA-B*40:01/Nuc_322-331_ tetramer. Results are representative of 6 independent uninfected donors. *Bottom:* PBMCs from convalescent COVID-19 individuals from the CHIRP cohort were analyzed by FACS for binding to the HLA-B*40:01/Nuc_322-331_ tetramer. Participant PID4103, but not participants PID4134 and PID 4135, harbors cells binding to the tetramer. Numbers correspond to the percentage of cells within the gates. Results are gated on live, singlet CD3+CD8+ cells. **(B)** Timeline of clinical course of PID4103’s SARS-CoV-2 infection and sampling. Red indicates the dates of specific symptom initiation and resolution, blue the dates and results of SARS-CoV-2 PCR tests, and green the dates of blood draws. **(C)** A distinct population of Nuc_322-331_-specific CD8+ T cells is detected by CyTOF in PID4103 through dual tetramer staining. PBMCs from one uninfected individual and from PID4103 were stained with two sets of HLA-B*40:01/Nuc_322-331_ tetramers conjugated to different metal lanthalides, facilitating specific detection of Nuc_322-331_-specific CD8+ T cells. Numbers correspond to the percentage of cells within the gates. Results are gated on live, singlet CD3+CD8+ cells. **(D)** Nuc_322-331_ specific CD8+ T cells can be stimulated by the Nuc_322-331_ peptide. PBMCs from PID4103 were phenotyped by CyTOF at baseline, or following 4 hours of co-stimulation with *α*CD49d/CD28 antibody in the absence or presence of the Nuc_322-331_ peptide. Stimulations were conducted in the presence of brefeldin A to enable the detection of IFN*γ*. Numbers correspond to the percentage of cells within the gates. Results are gated on live, singlet CD3+CD8+ cells.

We decided to focus our study on this participant for three reasons: 1) the presence of an immunodominant response detectable in her blood allowed us to have a sufficient number of cells to perform in-depth CyTOF phenotyping without combining multiple epitope specificities or pre-enrichment for tetramer+ cells; 2) the conservation of the immunodominant epitope among the variants-of-concern ensured that the memory response we studied would be relevant against the common globally circulating pathogenic strains, and 3) the patient reported symptoms of typical mild COVID-19 disease (as detailed below), and therefore can serve as a model for the typical course of disease experienced by most individuals who become infected with SARS-CoV-2 (19, 20).

Participant PID4103 is a 42-year-old Caucasian female whose course of SARS-CoV-2 infection has been extensively characterized (Fig. 1B, Table S2). The participant began experiencing a constellation of mild acute symptoms on 3/13/2020, including a fever of 102°F, sore throat, cough, chest pressure, and fatigue. Fever, cough, and sore throat resolved 2 days later, while chest pressure resolved 7 days post-symptom onset. Nine days post-symptom onset, she developed dysosmia/dysgeusia and headache, which lasted approximately one week. Complete resolution of all symptoms did not occur until 23 days from initial symptom onset. The participant tested positive by PCR for SARS-CoV-2 27 days post-symptom onset, and her clinical PCR result was confirmed by nasopharyngeal swab PCR at her baseline study visit 46 days post-symptom onset (407.5 and 161.5 copies/mL for N1 and N2 probes, respectively), which corresponded to 19 days after her first positive PCR test. The participant then attended follow-up visits at 1, 3, 10, and 23 weeks after her baseline visit. At all the follow-up study visits, she tested negative by PCR for the virus, in specimens from nasopharyngeal swabs, blood, stool, and urine. PID4103 reported no limitations to her activities of daily living over the course of disease.

She had no significant comorbidities other than a prior history of anxiety disorder and hypothyroidism for which she had previously received pharmacology therapy, and she had no concomitant medications during the study period. Her clinical lab tests by the time of her baseline visit (46 days post-symptom onset) were within normal limits. However, her ferritin levels, which have previously been shown to strongly correlate with COVID-19 symptoms (21), showed a downtrend over her five study visits (134, 120, 64, 56, 54 ng/mL). Her high sensitivity C reactive protein (hs-CRP) and erythrocyte sedimentation rate (ESR) levels were normal by the time she was enrolled in the study (<8.1 mg/L and <20 mm/hr, respectively), suggesting lack of systemic inflammation at any of the study visits. PID4103 reported no history of travel in the prior year, or past history of travel that may have coincided with exposure to SARS-CoV-1 (e.g., no 2002 travel to Canada or Asia). All together, these clinical features suggest PID4103 to have exhibited a typical case of mild COVID-19 that resolved on its own without medical intervention, and that did not result in any long-hauler symptoms.

### CyTOF characterization of Nuc_322-331_-specific CD8+ T cells through tetramer staining and peptide stimulation

To enable a deep assessment of the phenotypes of Nuc_322-331_-specific CD8+ T cells, we generated lanthanide-conjugated Nuc_322-331_ tetramers. This allowed for characterization of Nuc_322- 331_-specific CD8+ T cells by CyTOF, a technique that quantitates up to ∼40 proteins simultaneously at the single-cell level through mass spectrometric detection of metal-conjugated antibodies (22). To increase the noise-to-signal ratio and improve the specificity of detecting Nuc_322-331_-specific CD8+ T cells, we conjugated the same tetramer to two different lanthanides, and considered only cells binding both sets of tetramers as true positives. A population of Nuc_322- 331_-specific CD8+ T cells could be detected from PID4103 that was absent from uninfected individuals (Fig. 1C). To assess effector function, CD8+ T cells from PID4103 were examined by CyTOF in the absence of any stimulation, in the presence of costimulation with anti-CD49d/CD28 for 4 hours, and in the presence of costimulation with Nuc_322-331_ peptide, under conditions that enabled detection of peptide-induced cytokines at the single-cell level (5). Only in peptide-stimulated samples did we observe a distinct population of IFN*γ*-producing cells (Fig. 1D). These results collectively validate our ability to characterize Nuc_322-331_-specific CD8+ T cells by CyTOF, through tetramer staining as well as through identification of Nuc_322-331_-specific CD8+ T cells responding to cognate peptide stimulation.

### Longitudinal assessment Nuc_322-331_-specific CD8+ T cells reveals coordination with other components of antigen-specific adaptive immunity

Having validated the specificity of our reagents, we then characterized by CyTOF the magnitude of the Nuc_322-331_-specific CD8+ T cell response over the course of convalescence using the PBMCs isolated from PID4103 over the 5 study visits (Fig. 1B). The magnitude of the response over time was monitored by quantitating both the frequencies of tetramer+ cells in unstimulated specimens (Fig. 2A) and the frequencies of cells inducing IFN*γ* in response to Nuc_322-331_ stimulation (Fig. 2B). Both approaches enabled detection of Nuc_322-331_-specific CD8+ T cells at all 5 timepoints including the final one, which was >6 months after symptom onset. The overall phenotypes of the Nuc_322-331_-specific CD8+ T cells in the tetramer+ vs. IFN*γ*+ cells differed as visualized by tSNE (Fig. 2C). This observation is not surprising since the tetramer+ cells are not stimulated, while the IFN*γ*+ cells are. However, the majority of individually measured CyTOF markers were similar between the tetramer+ and IFN*γ*+ cells, and only IFN*γ* and TNF*α* appeared upregulated on the IFN*γ*+ cells relative to the tetramer+ cells (Fig. S2, S3). The latter is consistent with the need for *ex vivo* stimulation to detect cytokine production. Although there were more tetramer+ cells than IFN*γ*+ cells, their kinetics were similar over the time course, with a peak 67 days post-symptom onset (Fig. 2D). Our observation that there were approximately twice as many tetramer+ cells as IFN*γ*+ cells suggests that approximately half the CD8+ T cells of a given specificity are not captured by the IFN*γ* detection method. This was confirmed by examining the tetramer+ cells within the Nuc_322-331_-stimulated sample, half of which turned out to be IFN*γ*+ (Fig. S4). Interestingly, the majority of these IFN*γ*+ cells also produced TNF*α* (Fig. S4), suggesting a polyfunctional response. Importantly, at its maximal peak, the frequency of the Nuc_322-331_-specific CD8+ T cells as determined by tetramer staining was 1.3 x 10_-3_ (Fig. 2A), confirming the immunodominance of this epitope since COVID-19 CD8+ T cells with the most dominant epitope reported to date were detected at an average frequency of 6.88 x 10_-4_ (18).

**Figure 2.**
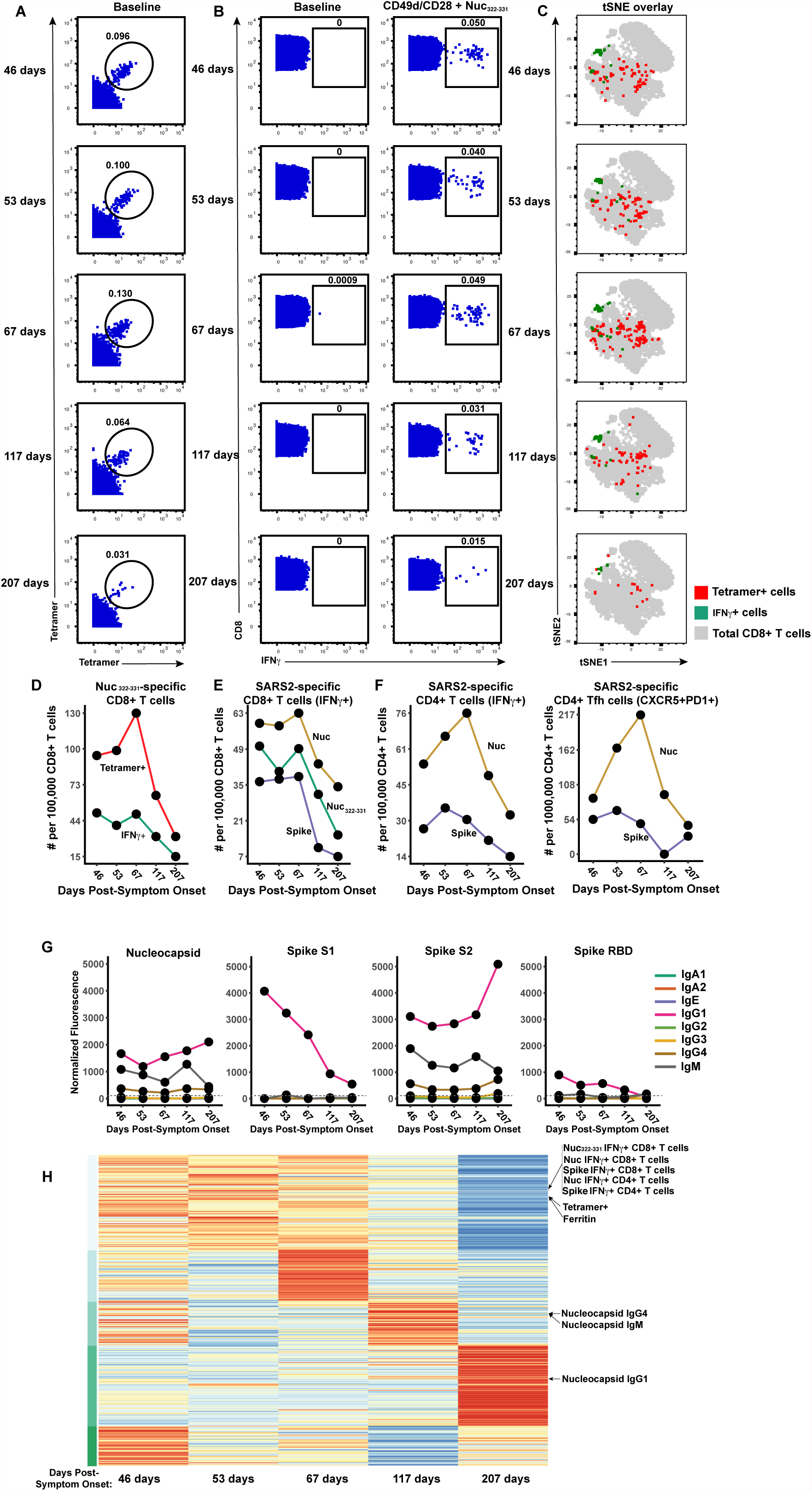
Longitudinal assessment of Nuc_322-331_-specific CD8+ T cell responses in PID4103 reveals coordination with other components of antigen-specific adaptive immunity. **(A)** Identification of Nuc_322-331-_specific CD8+ T cells by CyTOF. Baseline specimens that never underwent any stimulation were stained with HLA-B*40:01/Nuc_322-331_ tetramers detectable on two different CyTOF channels. The timeline refers to days since symptom onset. Numbers correspond to the percentage of cells within the gate. Results are gated on live, singlet CD3+CD8+ cells. **(B)** CD8+ T cells specifically producing IFN*γ* in response to Nuc_322-331_ stimulation were detected at all five timepoints. Numbers correspond to the percentage of cells within the gate. Results are gated on live, singlet CD3+CD8+ cells. **(C)** Tetramer+ and IFN*γ*+ cells responding to Nuc_322-331_ treatment reside in unique regions of the tSNE, suggesting phenotypic changes elicited by cognate peptide recognition. tSNE plots of total CD8+ T cells (*grey*), tetramer+ (*red*) from the baseline samples, and IFN*γ*+ (*green*) cells from the peptide-stimulated samples, over the course of convalescence of PID4103. Datasets correspond to those extracted from the data presented in *panels A* and *B*. **(D)** The tetramer+ response is higher in magnitude than the IFN*γ*+ response but exhibits similar kinetics, peaking 67 days post-symptom onset. Datasets correspond to those extracted from the data presented in *panels A* and *B*. **(E)** The responses of CD8+ T cell to Nuc_322-331,_ the entire nucleocapsid protein (Nuc), and the entire spike protein are coordinated. Note that the IFN*γ*+ response to Nuc_322-331_ is greater than the response to the entire spike proteins and lesser than the response to the entire nucleocapsid protein. **(F)** The total and Tfh CD4+ T cell responses against nucleocapsid peaks 67 days post- symptom onset, while the response to spike peaks slightly earlier. Total (*left*) or Tfh (CD4+CD45RO+CD45RA-PD1+CXCR5+) (*right*) CD4+ T cells responding to overlapping peptides spanning the entire nucleocapsid or spike proteins were assessed. **(G)** Titers of different antibody types against nucleocapsid, and the S1, S2, and RBD domains of spike, monitored at the 5 timepoints, and expressed as normalized fluorescence values (see Methods). The dotted line indicates the limit of detection. **(H)** Unsupervised k-means clustering of cells, antibodies and other biomarkers based on their abundance in PID4103’s blood across five time points. For each biomarker, abundance is normalized across time points and colored from red (highest) to blue (lowest). The CD4+ and CD8+ T cell against Nuc_322-331_, nucleocapsid, and spike clustered together. Interestingly, ferritin levels clustered close to them. In contrast, antibody responses against nucleocapsid were delayed and occurred after the peaks of the T cell responses. The green bars on the left correspond to clustering as determine by k-means.

We then assessed what fraction of the total nucleocapsid-specific CD8+ T cells is accounted for by Nuc_322-331_-specific cells. To this end, we stimulated samples from all five timepoints with overlapping peptides covering the entire nucleocapsid protein (Fig. S5A), and measured IFN*γ*-responding cells by CyTOF. Both cell populations (Nuc_322-331_ and total nucleocapsid-specific CD8+ T cells) had similar kinetics, as expected (Fig. 2E), and at all 5 timepoints, the Nuc_322-331_-specific CD8+ T cells accounted for the bulk of the nucleocapsid-specific CD8+ T cells (Fig. 2E). For comparison, we also assessed the spike-specific CD8+ T cells from these specimens using overlapping peptides covering the entire spike protein (Fig. S5A). These cells also mirrored the kinetics of the Nuc_322-331_-specific cells, but interestingly were always less abundant (Fig. 2E). These data reaffirm the immunodominance of Nuc_322-331_-specific CD8+ T cells, which surpass even that of the spike-specific CD8+ T cells. Furthermore, they demonstrate that the kinetics of the CD8+ T cell response against nucleocapsid and spike in PID4103 is coordinated, peaking 67 days post-symptom onset and decreasing thereafter.

We then assessed to what extent the Nuc_322-331_-specific CD8+ T cell response is coordinated with the response of CD4+ T cells and antibodies. When we assessed by CyTOF total and T follicular helper (Tfh) CD4+ T cells specific to nucleocapsid (Fig. 2F, S5B), we found that both peaked at the third timepoint (67 days post symptom onset), just like the Nuc_322-331_- specific CD8+ T cells did (Fig. 2E, F). When we measured 8 isotypes (IgM, IgG1, IgG2, IgG3, IgG4, IgA1, IgA2, and IgE) of antibodies against full-length nucleocapsid, we found that only IgM, IgG1, and IgG4 nucleocapsid-specific antibodies above detectable limits, with IgG1 being the dominant response. All three isotypes of nucleocapsid-specific antibodies increased from the third to the fourth timepoints (Fig. 2G). As a “helped” antibody response would be expected to develop only after a spike in a CD4+ Tfh response (which occurred at the third timepoint, Fig. 2F), these results suggest a coordinated T cell-dependent antibody response in this individual during convalescence. For comparison, we also assessed the CD4+ T cell and antibody response against spike. Spike-specific CD4+ T cells peaked slightly earlier than the Nuc_322-331_-specific CD8+ T cells (Fig. 2F). Antibodies against the spike N-terminal S1 and C-terminal S2 domains, as well as against the RBD domain of S1 that mediates binding to the ACE2 entry receptor, were quantitated. Interestingly, in contrast to the T cell data where the response to nucleocapsid response dominated over the response to spike, the antibody response to spike was dominant over the response to nucleocapsid (Fig. 2G). Similar to the nucleocapsid data, the dominant antibody isotype against spike was IgG1. Interestingly, however, the antibody response to S1 and RBD progressively decreased over the course of convalescence, while the antibody response to S2 more closely mirrored the response to nucleocapsid, increasing from the third to the fourth timepoints (Fig. 2G). Taken together, these data suggest a synchronous increase in CD4+ and CD8+ T cells preceding the antibody response.

As a complementary way of examining the coordination between these different adaptive immune responses, we conducted an integrated analysis of all the SARS-CoV-2-specific T cell and antibody response measurements from our study. We included the frequencies of all the subsets of Nuc_322-331_-, and Nuc-, and Spike-specific T cells we identified by manual gating as well as flowSOM clustering (see subsequent sections); the median expression levels of each CyTOF- quantitated antigen on total CD8+ T cells, the tetramer+ cells, and all the CD4+ and CD8+ IFN*γ*+ cells responding to Nuc_322-331_, nucleocapsid, or spike peptide treatments; all 32 antibody measurements (4 proteins x 8 isotypes); and the clinical lab measurements. This resulted in a matrix of 393 measured parameters, for each of the 5 timepoints. When we conducted k-means unsupervised clustering to assess which parameters were closely related, we found the kinetic patterns of the CD4+ and CD8+ T cell responses to Nuc_322-331_, nucleocapsid, and spike to cluster together (Fig. 2H). Interestingly, the levels of ferritin, which were reported to positively correlate with COVID-19 symptoms (21), clustered right next to the magnitude of the tetramer+ cell response (Fig. 2H). As expected, the nucleocapsid antibody response clustered separately as it was delayed relative to the T cell response (Fig. 2H). Overall, these data suggest that the Nuc_322- 331_-specific CD8+ T cell response is synchronized with CD4+ and CD8+ T cell responses against nucleocapsid and spike, followed by a boosting of the nucleocapsid and S2 IgG1, IgG4, and IgM antibodies.

### The phenotypes and potential for long-term persistence of Nuc_322-331_-specific CD8+ T cells evolve during convalescence

We then took advantage of the 38-parameter phenotyping of our tetramer+ cells to characterize the phenotypes of Nuc_322-331_-specific CD8+ T cells. CD8+ T central memory (Tcm), T effector memory (Tem), T transitional memory (Ttm), Temra, and a mixed population of naïve and T stem cell memory cells (Tn/Tscm) were identified through use of various combinations of the phenotyping markers CD8, CD45RO, CD45RA, CCR7, and CD27 (Fig. 3A). In addition we distinguished, among the memory (CD45RO+) CD8+ T cells, those that were less (CD27+CD28+) or more (CD27-CD28-) terminally differentiated (Fig. 3A). Terminal differentiation and expansion potential were also examined by monitoring expression of CD57 (terminal differentiation marker), CD27 (marker of long-lived cells), and CD127 (alpha chain of IL7 receptor involved in homeostatic proliferation) (Fig. 3B). Cytolytic activity was assessed by monitoring expression of the effector molecules perforin and granzyme B, the degranulation marker CD107a, and CD29, which marks cells with cytolytic activity (23) (Fig. 3C). Tetramer+ cells could be detected among all the aforementioned populations, although in vastly different proportions (Fig. 3A-C).

**Figure 3.**
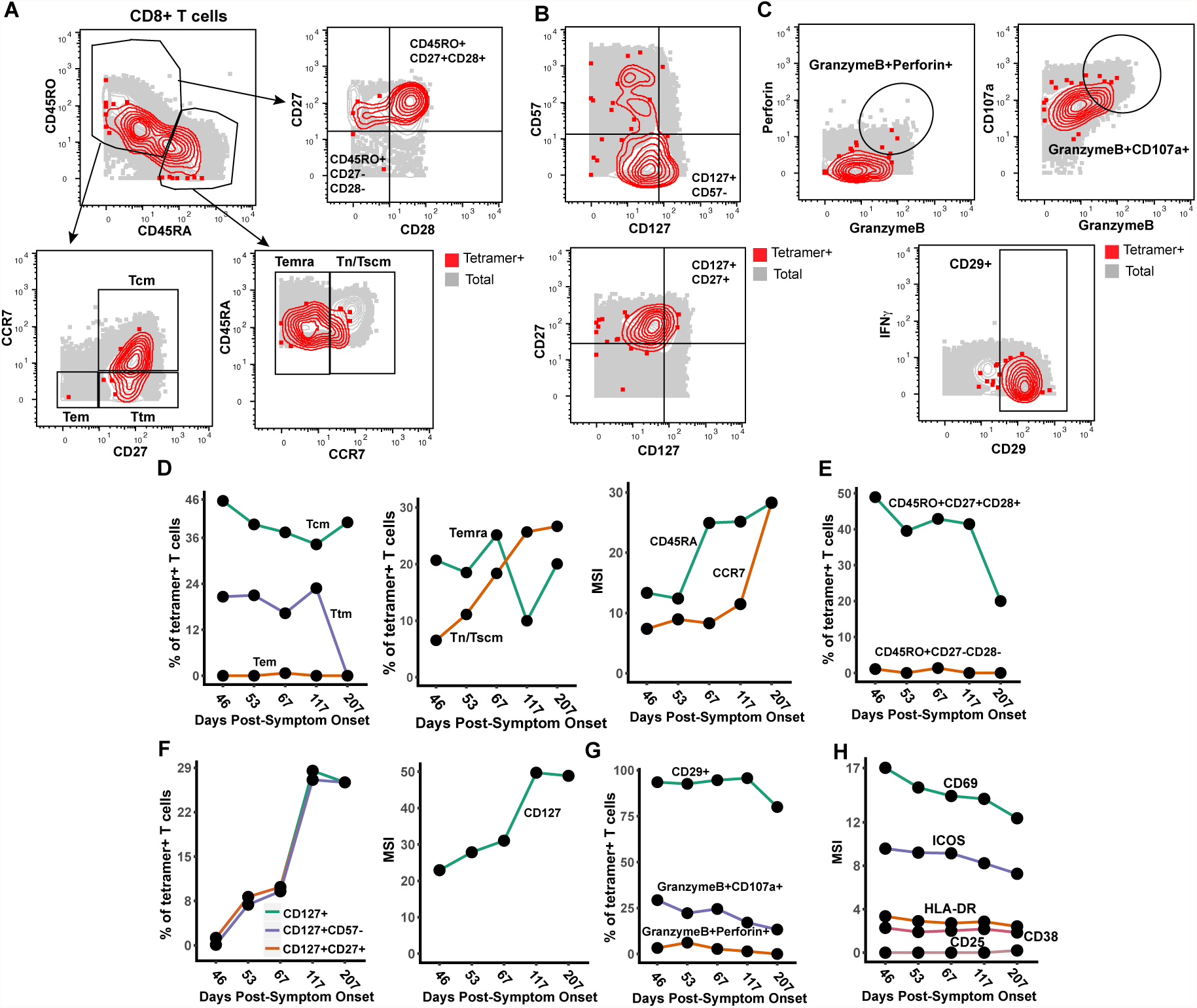
Nuc_322-331_-specific CD8+ T cells in PID4103 slowly differentiate over the course of convalescence to a less activated state more capable of expanding and migrating to lymph nodes. **(A)** Gating strategy to identify CD8+ Tcm, Tem, Ttm, Temra, Tn/Tscm at early and late differentiation stages. The Nuc_322-331_-specific CD8+ T cells (tetramer+) cells are shown as red contours, while total CD8+ T cells are shown in grey. The following gates were used: Tcm (CD45RO+CD45RA-CD27+CCR7+), Ttm (CD45RO+CD45RA-CD27+CCR7-), Tem (CD45RO+CD45RA-CD27-CCR7-), Temra (CD45RO-CD45RA+CCR7-), Tn/Tscm (CD45RO-CD45RA+CCR7+), early differentiation (CD45RO+CD45RA- CD27+CD28+) and late differentiation (CD45RO+CD45RA-CD27-CD28-). **(B)** Gating strategy to identify different populations of CD127+ cells among total (*grey*) and Nuc_322-331_-specific (*red*) CD8+ T cells. Shown also are gates for less differentiated (CD57-) and Tcm-like (CD27+) CD127+ T cells. **(C)** Gating strategy to identify cytolytic Nuc_322-331_-specific CD8+ T cells. *Top:* Gates defining CD8+ T cells co-expressing granzyme B and perforin, or granzyme and CD107a. *Bottom:* Gate defining cells expressing high levels of CD29, a marker for cytolytic CD8+ T cells. **(D)** The proportions of tetramer+ cells belonging to the Tcm, Tem, Ttm, Temra, and Tn/Tscm subsets as defined in *panel A* are shown in the first two plots. Note the high contribution of Tcm at all timepoints, and the progressive increase of the Tn/Tscm subset over time. The panel on the right displays the median expression levels of CD45RA and CCR7 (markers used to define the Tn/Tscm subset [*panel A*]) within the tetramer+ population. **(E)** Early differentiated CD8+ T cells steeply decline in abundance only at the final timepoint, 207 days post symptom onset. Shown are the proportions of tetramer+ cells belonging to the early (CD45RO+CD27+CD28+) and late (CD45RO+CD27-CD28-) differentiated subsets over the course of convalescence. **(F)** Progressive increase in CD127+ Nuc_322-331_-specific CD8+ T cells over a ∼6 month period of convalescence. *Left:* Proportions of tetramer+ cells that were CD127+, CD127+CD57- and CD127+CD27+. The overlapping frequencies of the three populations of cells suggest that most of the CD127+ cells are CD57- and CD27+. *Right:* Median expression levels of CD127 within the tetramer+ population. **(G)** Cytolytic Nuc_322-331_-specific CD8+ T cells slowly decrease over the course of convalescence. The proportions of tetramer+ cells that were CD29+, granzymeB+CD107a+, and granzymeB+perforin+ are shown. **(H)** The activation state of Nuc_322-331_-specific CD8+ T cells generally decreases slowly over the course of convalescence. The median expression levels of the indicated activation markers on tetramer+ cells are shown. A gradual decrease was apparent among CD69, ICOS, HLADR, CD38 but not CD25, whose expression was low at all timepoints.

We assessed the tetramer+ cells for relative changes in subset distribution over the ∼6- month period analyzed in this study. Among tetramer+ cells, Tem cells were at negligible frequencies throughout the timecourse, while Tcm cells were most common. Tetramer+ Ttm and Temra cells were also abundant, although the Ttm subset frequency dropped precipitously at the final timepoint. Interestingly, the contribution of the Tn/Tscm subset increased steadily over the course of convalescence, reaching the highest levels at the final timepoint (Fig. 3D). This increase parallels the increase in expression levels of CD45RA and CCR7 (markers used to define the Tn/Tscm subset) within the tetramer+ population (Fig. 3D). In terms of differentiation state, there was a progressive decrease over time of the early-differentiated CD27+CD28+ memory T cell subset among tetramer+ cells (Fig. 3E). This was accompanied by a progressive increase in CD127 positivity, with the CD127+ cells residing almost exclusively among CD57- and CD27+ cells (Fig. 3F). Cytolytic tetramer+ cells decreased over time (Fig. 3G), and this was accompanied by a gradual decrease in the expression levels of some (CD69, ICOS, HLADR, CD38) but not all (CD25) activation markers on these cells (Fig. 3H). Together, these results suggest a continual differentiation of Nuc_322-331_-specific CD8+ T cells long after resolution of infection. These changes include the evolution of the cells to a state defined by less activation and cytolytic activity, and more proliferative and expansion potential.

### Clustering of high-dimensional datasets identifies features of convalescence-associated expanding cluster of Nuc_322-331_-specific CD8+ T cells

The subset identification based on manual gating described above utilizes only a small fraction of the phenotyping markers examined by our CyTOF panel. Visualization of the phenotypic distribution of the tetramer+ cells by tSNE suggests global changes in phenotypes over time (Fig. 4A) that may not have been captured through manual gating. We therefore next implemented a more holistic approach of subset definition by clustering. Total CD8+ T cells in the unstimulated specimens (including the tetramer+ cells) were segregated into 5 clusters by flowSOM (24) (Fig. 4B). Although tetramer+ cells were detected among all 5 clusters, they were not evenly distributed, and the distribution also changed over time (Fig. 4C, D). Clusters A2 and A4 were the dominant clusters among tetramer+ cells, but interestingly while cluster A2 increased over time, cluster A4 decreased (Fig. 4D). When we manually gated on a concatenated file corresponding to all of the cluster A2 and A4 cells (which together include most of the tetramer+ cells), we found that these cells collectively harbored Tcm, Tem, Ttm, Temra, and Tn/Tscm subsets (Fig. 4E), suggesting that the dominant population of tetramer+ cells cannot be binned into any single canonical cellular subset. To try to define features of clusters A2+A4, we assessed for antigens that were similarly differentially expressed on these two clusters as compared to total CD8+ T cells. Relative to total CD8+ T cells, A2 and A4 cells expressed high levels of CD127 and the transcription factor NFAT1, as well as high levels of the lung-homing molecules CD49d, CD29, and CCR5 (Fig. 4F). We then assessed for differentially-expressed markers that were unique to clusters A2 vs. A4, to assess the subset features that increase (cluster A2) vs. decrease (cluster A4) over the course of convalescence. Cluster A2 expressed high levels of the lymph node homing receptors CCR7 and CD62L, the checkpoint molecules TIGIT and CTLA-4, the co-stimulatory receptors CD28 and Ox40, and the anti-apoptotic protein BIRC5 (Fig. 4G). In comparison, cluster A4 expressed low levels of CCR7 and CD62L, high levels of the activation marker CD69, and high levels of the degranulation marker CD107a (Fig. 4H). These data are consistent with the manual gating results suggesting a slow expansion of long-lived Nuc_322-331_- specific CD8+ T cells paired with a decrease in their cytolytic counterparts, but identify additional phenotypic features of the cellular subsets to which these cells belong.

**Figure 4.**
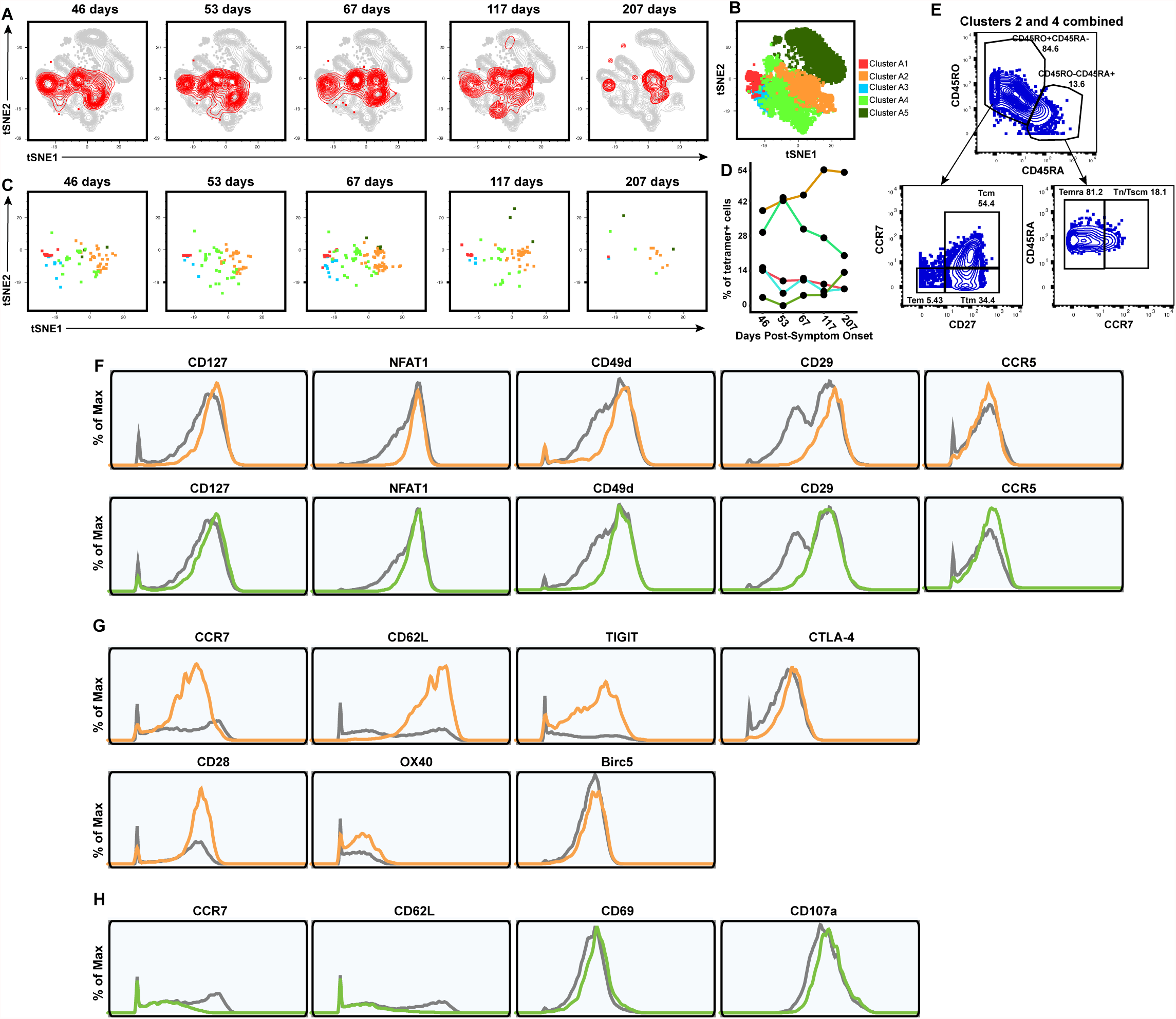
Clusters of Nuc_322-331_-specific CD8+ T cells from PID4103 exhibit different expansion and contraction. **(A)** The overall phenotypes of Nuc_322-331_-specific CD8+ T cells change over the course of convalescence. tSNE plots of total (*grey*) and tetramer+ (*red*) CD8+ T cells as a function of time since symptom onset. **(B)** FlowSOM clusters of CD8+ T cells. Total CD8+ T cells (including the tetramer+ cells) were clustered by flowSOM to identify five clusters. The location of each cluster is mapped onto the tSNE space depicted in *panel A*. **(C)** Distribution over time of Nuc_322-331_-specific CD8+ T cells among the five clusters identified in *panel B*. **(D)** Proportion of Nuc_322-331_-specific CD8+ T cells in each cluster as a function of time since symptom onset. The dominant clusters, A2 and A4, increase and decrease over time, respectively. **(E)** Clusters A2 and A4 include multiple cellular subsets. Gating strategy showing the identification of the Tcm, Tem, Ttm, Temra, and Tn/Tscm subsets, all of which were well-represented among the two dominant clusters. **(F)** Phenotypic features shared by clusters A2 and A4. Relative to total CD8+ T cells, clusters A2 and A4 expressed high levels of CD127, the transcription factor NFAT1, and the lung homing receptors CD49d, CD29, and CCR5. Total CD8+ T cells are depicted in grey, cluster A2 cells in orange, and cluster A4 cells in green. **(G)** Phenotypic features exhibited by cluster A2 and not A4. Cluster A2, whose contribution among tetramer+ cells increased over the course of convalescence, expressed high levels of the lymph node homing receptors CCR7 and CD62L, the checkpoint molecules TIGIT and CTLA4, the co-stimulatory molecules CD28 and Ox40, and the pro-survival factor BIRC5. Total CD8+ T cells are depicted in grey and cluster A2 cells in orange. **(H)** Phenotypic features exhibited by cluster A4 and not A2. Cluster A4, whose contribution among tetramer+ cells decreased over the course of convalescence, expressed low levels of the lymph node homing receptors CCR7 and CD62L, high levels of the activation marker CD69, and high levels of the degranulation marker CD107a.

### Polyfunctional Nuc_322-331_-specific CD8+ T cells are detected months into convalescence

While phenotypic analysis of the tetramer+ cells in the unstimulated samples enabled an in-depth assessment of the differentiation states, expansion potential, homing patterns, and cytolytic activity of Nuc_322-331_-specific CD8+ T cells, they did not allow assessment of the cytokines these cells are capable of producing. We therefore next characterized the phenotypes of cells from the specimens stimulated for 4 hours with Nuc_322-331_ peptide. In these specimens, Nuc_322-331_- specific CD8+ T cells were defined as the IFN*γ*+ cells following peptide treatment (Fig. 2B, D), similar to recently implemented methods (5). Characterization of the canonical subsets (Tcm, Tem, Ttm, Temra, Tn/Tscm) revealed that in contrast to the data from unstimulated tetramer+ cells (Fig. 3D), Tcm cells were not the dominant subset among the responding cells (Fig. 5A, C). This can be explained by the decreased expression of the Tcm markers CD45RO and CCR7 on the IFN*γ*+ as compared to the tetramer+ cells (Fig. S2, S3), likely caused by stimulation-induced downregulation of these receptors. Similar to the tetramer+ data, however, the contribution of the Tn/Tscm subset to the IFN*γ*+ cells increased over time (Fig. 5C). Cytolytic cells were detected among the IFN*γ*+ cells (Fig. 5D). Although these IFN*γ*+ cytolytic cells decreased over time, they still represented a sizable proportion of the cells up until the 4_th_ timepoint, suggesting the existence of polyfunctional (both IFN*γ*-producing and cytolytic) Nuc_322-331_-specific CD8+ T cells well into convalescence (at least 117 days post-symptom onset). To further probe the polyfunctionality of Nuc_322-331_-specific CD8+ T cells, we assessed to what extent they induced IFN*γ*, TNF*α*, or IL6, the latter of which has been implicated in COVID-19 pathogenesis and may be produced by multiple immune cells including T cells (25). At the first four timepoints, the vast majority of IFN*γ*+ cells also produced TNF*α*, while at the final timepoint responding cells were more equally distributed among IFN*γ*+TNF*α*+ and IFN*γ*+TNF*α*- cells (Fig. 5E, 5F). Interestingly, none of the IFN*γ*+ cells co-produced IL6, and overall the proportion of Nuc_322-331_-specific IL6+ T cells was negligible (Fig. 5E). Assessing all three cytokines together, it was apparent that most of the responding cells dually produced IFN*γ* and TNF*α* (but no IL6), and that the frequencies of these cells decreased over time beginning at the third timepoint, 67 days post-symptom onset (Fig. 5F). Not only did the percent of IFN*γ*+TNF*α*+ decrease over time, but also the absolute levels of IFN*γ* and TNF*α* produced by these cells (Fig. 5G).

**Figure 5.**
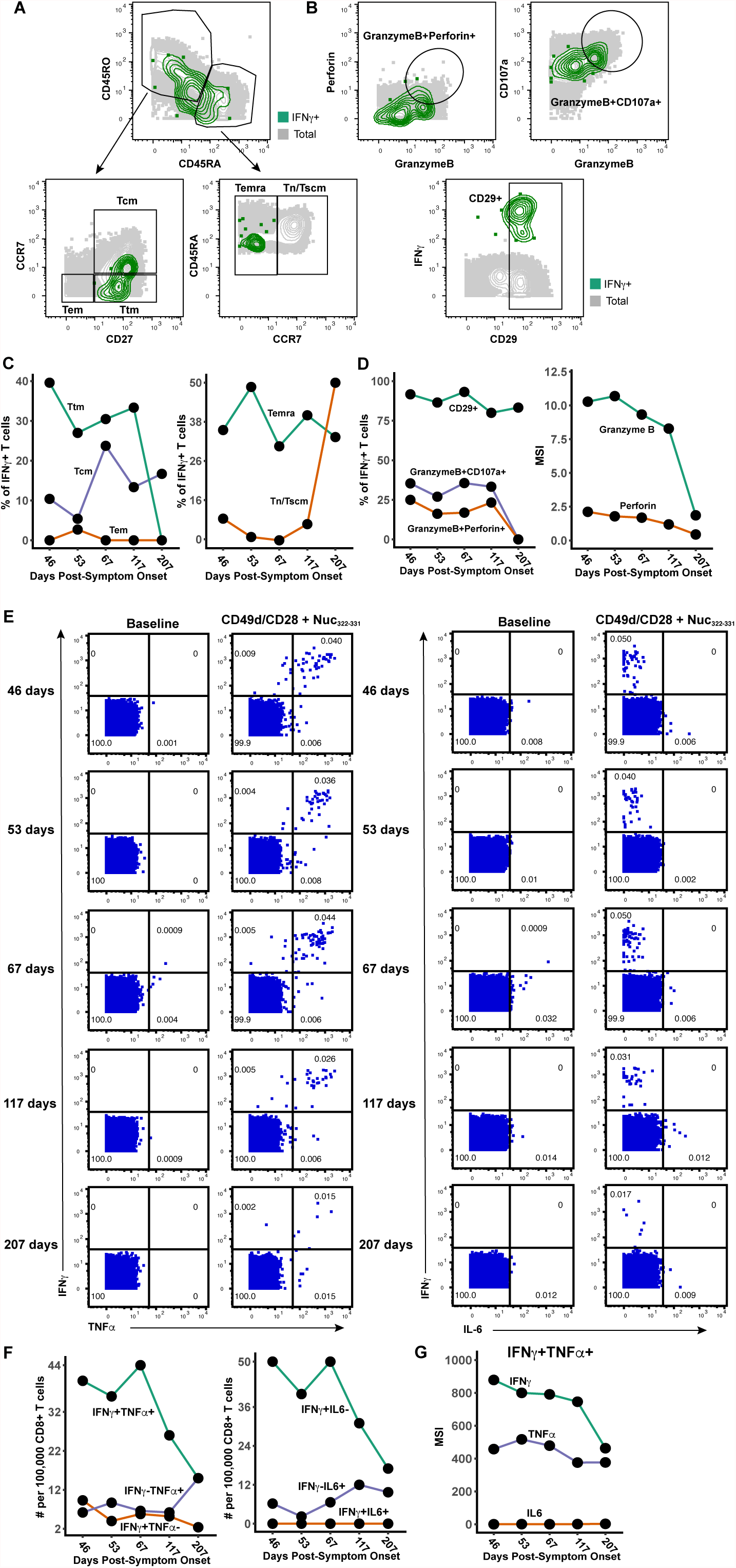
Polyfunctional Nuc_322-331_-specific CD8+ T cells are detected months into PID4103’s convalescence. **(A)** Gating strategy to identify the Tcm, Tem, Ttm, Temra, and Tn/Tscm subsets of Nuc_322- 331_-specific CD8+ T cells, identified as those responding to peptide stimulation by producing IFN*γ*. The Nuc_322-331_-specific CD8+ T cells (IFN*γ*+) cells are shown as green contours, while total CD8+ T cells are shown in grey. Subset definitions are identical to those used in Figure 3A. **(B)** Gating strategy to identify cytolytic Nuc_322-331_-specific CD8+ T cells among those inducing IFN*γ* upon cognate peptide stimulation. *Top:* Gates defining CD8+ T cells co- expressing granzyme B and perforin, or granzyme and CD107a, are indicated. *Bottom:* Gate defining cells expressing high levels of CD29, a marker for cytolytic CD8+ T cells. **(C)** Proportion of IFN*γ*+ Nuc_322-331_-specific CD8+ T cells belonging to the Tcm, Tem, Ttm, Temra, and Tn/Tscm subsets as defined in *panel A*. The lower contribution of Tcm at all timepoints is likely mediated by activation-induced CCR7 downregulation. Similar to the tetramer data (Fig. 3D), an increase in the contribution of the Tn/Tscm subset was observed over time. **(D)** Cytolytic Nuc_322-331_-specific CD8+ T cells slowly decrease over the course of convalescence. *Left:* Proportion of IFN*γ*+ cells that were CD29+, granzymeB+CD107a+, and granzymeB+perforin+. *Right:* Median expression levels of the indicated cytolytic activity markers on the IFN*γ*+ cells. **(E)** Most Nuc_322-331_-specific CD8+ T cells responding to peptide stimulation secrete multiple cytokines. Dot plots showing the expression of IFN*γ* and TNF*α* (*left*) or IFN*γ* and IL6 (*right*) on CD8+ T cells among baseline or peptide-stimulated samples. Numbers correspond to the percentage of cells within the gates. Results are gated on live, singlet CD3+CD8+ cells. Most responding Nuc_322-331_-specific CD8+ T cells were IFN*γ*+TNF*α*+IL6-. **(F)** The proportion of IFN*γ*+TNF*α*+IL6- CD8+ T cells responding to Nuc_322-331_ stimulation decreases over the course of convalescence. The cell populations are taken from the gates shown in *panel E*. **(G)** The level of IFN*γ* and TNF*α* produced by Nuc_322-331_-specific CD8+ T cells decreases over the course of convalescence, as shown by median signal intensity of the IFN*γ*+TNF*α*+ cells.

Finally, to further probe the effector features of these polyfunctional responding cells, we implemented flowSOM clustering. As we had conducted CyTOF phenotyping of CD8+ T cells responding not only to Nuc_322-331_, but also to overlapping peptides from the entire nucleocapsid and spike proteins (Fig. 2E), we clustered all of these responding cells together to compare their effector functions and overall phenotypes. The overall phenotypes of the Nuc_322-331_-specific CD8+ T cells were similar to those of the nucleocapsid-specific CD8+ T cells (Fig. 6A), consistent with the response to this immunodominant peptide being representative of the response to its parent protein. Interestingly, however, the phenotypes of the spike-specific CD8+ T cell cells differed as reflected by their different distribution within the tSNE plot (Fig. 6A). Furthermore, while the majority of the Nuc_322-331_-specific and Nuc-specific CD8+ T cells belonged to cluster B1, the phenotypes of the spike-specific CD8+ T cells were more evenly distributed, although cluster B1 was also the dominant cluster for the spike-specific cells (Fig. 6B). Cluster B1 included T cells expressing cytolytic markers (granzyme B, CD107a) and cytokines (IFN*γ*, TNF*α*) (Fig. 6C) consistent with the polyfunctionality of SARS-CoV-2-specific CD8+ T cells during convalescence. Cluster B1, however, also included cells with low levels of cytolytic effectors, and these became more prevalent at the later timepoints (Fig. 6D), consistent with the diminishing polyfunctionality of SARS-CoV-2-specific CD8+ T cells over the course of convalescence and their differentiation into long-lived memory cells.

**Figure 6.**
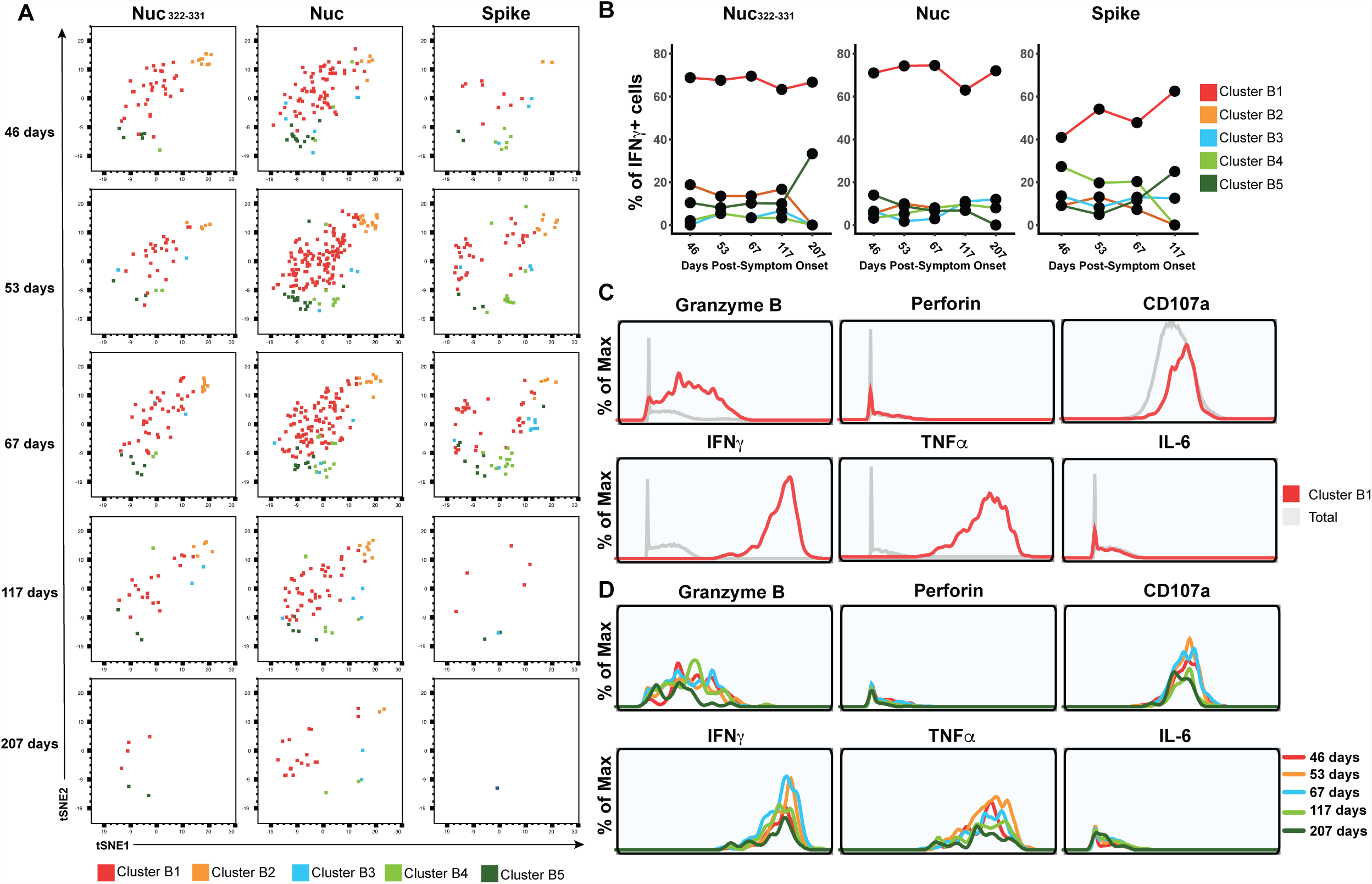
PID4103’s CD8+ T cells responding to Nuc_322-331_ stimulation are more similar to those responding to nucleocapsid than to spike peptides. **(A)** Cluster distribution of CD8+ T cells responding to Nuc_322-331_, or to peptides spanning the entire nucleocapsid or spike proteins. IFN*γ*+ CD8+ T cells from the Nuc_322-331_, nucleocapsid, or spike-stimulated specimens were split into five clusters (B1-B5) by flowSOM. The responding cells are shown as dot plots and colored according to their cluster membership. Note the higher similarity of cells in the tSNE among the Nuc322-331- and Nuc-specific cells, relative to the spike-specific ones. **(B)** Cluster B1 is dominant among CD8+ T cells with all three specificities, but more prominent among the Nuc_322-331_- and Nuc-specific cells. **(C)** Cluster B1 cells, to which most cells responding to Nuc_322-331_, Nuc, and Spike stimulation CD8+ T cells belong, are characterized by high expression levels of the cytolytic markers granzyme B and CD107a, and the cytokines IFN*γ* and TNF*α*. **(D)** The sub-populations of cluster B1 cells expressing higher levels of effector cytokines and cytolytic molecules decrease over the course of convalescence. Shown are histogram plots depicting cluster B1 cells colored according to timepoint. Although all the cells shown belong to cluster B1, those from the later timepoints expressed lower levels of granzyme B, CD107a, IFN*γ*, and TNF*α*.

## DISCUSSION

An important role for CD8+ T cells in effective host response against SARS-CoV-2 was implicated early during the COVID-19 pandemic, when it was observed that lymphopenia, particularly of CD8+ T cells, associates with disease severity (26). Moreover, the CD8+ T cell response directed against the nucleocapsid protein of SARS-CoV-2 may be particularly important, as it appears to be more common than the response directed against spike and other non-structural proteins (14, 17, 27). Interestingly, CD8+ T cell responses against SARS-CoV-1 also appear to be nucleocapsid-focused, and SARS-CoV-1-specific CD8+ T cells still detectable 17 years after the 2002 SARS outbreak were found to be reactive against nucleocapsid (7).

These prior observations implicate nucleocapsid-specific CD8+ T cells not only as important guardians during acute infection, but also as a reservoir of long-lived memory cells.

In this study, we screened both spike and nucleocapsid MHC class I tetramers against convalescent COVID-19 participants from the CHIRP cohort, and identified Nuc_322-331_ as an immunodominant epitope in one of the participants. The maximal proportion of Nuc_322-331_- specific CD8+ T cells was 0.13%, which exceeds the ∼0.0688% that was recently reported as the most immunodominant CD8+ T cell response known to date in COVID-19 (18) and is about two orders of magnitude higher than the proportion of SARS-CoV-2 CD8+ T cells against other epitopes (16). Further supporting its immunodominance is our observation that at all five timepoints, the proportion of Nuc_322-331_-specific cells exceeded that of the CD8+ T cells directed against the entire spike protein. Other groups have also reported Nuc_322-331_-specific CD8+ T cells at frequencies similar to ours (0.093%(13)) or lower (<0.01%; (17)). The lower frequency detected in the latter study may be donor-dependent, or could reflect a frequency calculation based off measurements of pre-enriched tetramer+ cells. Of note, the relatively high frequency of Nuc_322-331_-specific CD8+ T cells is not due to prior clonal amplification elicited by common-cold coronaviruses, as this peptide is not conserved in those strains. It is also unlikely to have been previously primed by SARS-CoV-1 in PID4103, since despite 100% conservation of this peptide between the two SARS strains, the participant’s travel history suggest that she could not have been previously exposed to the 2002 virus. Whether the immunodominance of Nuc_322-331_- specific CD8+ T cells is due to high frequencies of precursors in the naïve TCR repertoire, the molecular features of this peptide interacting with MHC, and/or the kinetics of its presentation by antigen presenting cells *in vivo*, remains to be determined. Importantly, as the Nuc_322-331_ sequence is 100% identical in all of the current SARS-CoV-2 variants of concern, the Nuc_322-331_ responses characterized in this study can be presumed to be as effective against the variants as they are against the original strain.

Extensive CyTOF phenotyping of tetramer+ CD8+ T cells of the longitudinal samples from PID4103 revealed that among the canonical subsets, Tcm were most common. These results are consistent with recent reports that SARS-CoV-2 nucleocapsid-derived tetramer+ cells were commonly of the CD45RA-CD27+ phenotype (18), characteristic of both the Tcm and Ttm subsets. We further observed that one of the surface markers used to define Tcm cells, CCR7, increased over the course of convalescence among tetramer+ cells. Furthermore, unbiased clustering of the datasets identified a dominant cluster (A2) of tetramer+ cells that increased over the course of convalescence, and this cluster was characterized by high levels of both CCR7 and CD62L expression. As both of these receptors direct immune cells into lymph nodes, the data suggest that over the course of ∼6 months of convalescence, Nuc_322-331_-specific CD8+ T cells continuously differentiate towards a state more likely to home to lymph nodes.

Tcm cells, a canonical lymph node-homing subset, are thought to be relatively long-lived compared to other memory subsets such as Tem and Ttm (28). Another long-lived subset expressing CCR7 are Tscm cells, which have long telomeres and are maintained by ongoing proliferation (29). Although in our CyTOF panel we could not distinguish between Tn and Tscm cells, we suspect that the CD45RA+CD45RO-CCR7+ subset (comprising both Tn and Tscm cells) we detected among tetramer+ cells were predominantly of the Tscm phenotype, since naïve T cells recognizing Nuc_322-331_ are expected to be very rare. These Tn/Tscm cells increased steadily over the course of convalescence, plateauing at 117 days post-symptom onset. A recent cross-sectional study where convalescents were binned into early vs. late convalescence also found CD45RA+CCR7+ tetramer+ cells to be higher in the latter phase (15), consistent with our longitudinal data. That study also found that among tetramer+ cells, the percentage of CD127+ cells increased as a function of days since symptom onset out to 120 days. This again mirrors our longitudinal analysis, where we found that both the percentage of CD127+ cells and the median expression levels of CD127 on the cells steadily increased from 43 to 207 days post symptom onset. Interestingly, the CD127+ cells were almost exclusively CD27+ and CD57-, suggesting a high expansion potential for these cells. Indeed, our recent study demonstrated that SARS-CoV-2-specific T cells expressing CD127 were capable of homeostatically expanding *ex vivo* (5). Together with prior demonstrations that CD127 expression identifies CD8+ T cell memory precursors giving rise to long-lived memory cells (30), these data suggest a slow differentiation of SARS-CoV-2-specific CD8+ T cells to lymph-node homing, long-lived memory cells with expansion potential, in the months following recovery from mild COVID-19.

In contrast, other features of the Nuc_322-331_-specific CD8+ T cells tended to decrease over time. Within the first four months of convalescence, Nuc_322-331_-specific CD8+ T cells were largely polyfunctional, with most producing both IFN*γ* and TNF*α*, and a substantial fraction of these cells additionally expressing cytolytic markers. This changed at the final timepoint (>6 months convalescence), when the cytolytic activity of the cells dropped and cells capable of producing both IFN*γ* and TNF*α* decreased. Interestingly, the decrease in polyfunctionality was mirrored by a decrease in the activation state of the cells, as reflected by a decrease in CD69 and ICOS, and to a lesser extent HLADR and CD38. This was in some aspect surprising as we would have expected the activation state of the cells to have returned back to normal by the second timepoint, which was almost 2 months after symptom onset and 30 days after the complete resolution of all symptoms. Together, these results suggest slow changes in the features and functional responses of SARS-CoV-2-specific CD8+ T cells long after full recovery from mild COVID-19.

One of the most interesting findings from this study was the coordinated response of Nuc_322-331_-specific CD8+ T cells with other components of the adaptive immune system. The magnitude of the Nuc_322-331_-specific CD8+ T cell response peaked at 67 days post symptom onset, and then decreased thereafter, as did the total CD4+ and CD8+ T cell response against nucleocapsid, and the total CD8+ T cell response against spike. The antibody response also exhibited convalescence spikes, but delayed relative to the T cell peak. The delay of the antibody response is reasonable given the time needed for B cells to be effectively helped by CD4+ T cells, and our finding that the nucleocapsid-specific CD4+ T cell response peaked at the 3_rd_ timepoint, which was followed by spikes at the 4_th_ timepoint in multiple isotypes of antibodies against the same protein, suggests a CD4+ T cell-helped B cell response in PID4103. An increase in the T cell and antibody response 67 to 117 days post symptom onset was in fact unexpected, given that this was 30-94 days post-symptom resolution and long after the virus had been cleared. Although this convalescence peak in adaptive immune response could theoretically have been caused by re-infection of this participant, we think that possibility unlikely because 1) she tested negative in PCR tests at all timepoints except her baseline visit, 2) we saw no evidence of elevated Nuc_322-331_-specific CD8+ T cell activation at the spiking timepoints (in fact activation progressively decreased, as discussed above), which we would have expected upon re-infection, 3) the participant reported no reemergence of COVID-19 symptoms after initial symptom resolution, 4) re-infection is overall uncommon. Further in-depth analyses of the SARS-CoV-2-specific immune responses in additional individuals are needed to confirm whether the immune response detailed here is recapitulated in others who experienced mild COVID-19 disease.

In summary, we report an unexpectedly dynamic evolution of Nuc_322-331_-specific CD8+ T cells during convalescence in PID4103. This evolution was gradual and persistent even up to 6 months after complete symptom resolution. We observed coordination of this response with the CD4+ T cell and antibody responses directed against the same antigen, and found that it was characterized by a progressive diminution of the activation state and polyfunctionality of cells in parallel with increases in their expansion potential. If one assumes similarities to nucleocapsid-specific CD8+ T cells from SARS-CoV-1, then the course of differentiation we describe here may be one that leads to SARS-CoV-2-specific memory CD8+ T cells that can persist for up to 17 years, and perhaps even longer.

## METHODS

### Human Subjects

This study was approved by the University of California, San Francisco (IRB #20-30588). Informed consent was obtained from all subjects. The study utilized specimens from the UCSF acute COVID-19 Host Immune Pathogenesis (CHIRP) longitudinal cohort. Five longitudinal specimens were collected from acute COVID-19-infected individuals, the first within 31 days of symptom onset or SARS-CoV-2 exposure (week 0, baseline visit), followed by collections at 1, 3, 10, and 24 weeks from baseline visit. Whole blood was collected in ethylenediamine tetraacetic acid (EDTA) tubes and peripheral blood mononuclear cells (PBMCs) were isolated by ficoll as previously described (5) and cryopreserved in 10% DMSO diluted in fetal bovine serum (FBS). Plasma from the same specimens were collected and cryopreserved. A total of 21 CHIRP participants with confirmed SARS-CoV-2 infection as assessed by RT-PCR were screened by FACS for specific binding to APC-labeled MHC class I tetramers, as described in the following section. PID4103 (Table S1) was identified as a donor with an immunodominant response against the HLA-B*40:01/Nuc_322-331_ tetramer.

### Flow Cytometry

Cryopreserved PBMCs from 21 CHIRP participants were thawed and cultured overnight to enable antigen recovery, and then screened by FACS for specific binding to APC-labeled MHC class I tetramers (Table S2) obtained from the NIH Tetramer Core Facility. These tetramers harbor predicted MHC class I epitopes from SARS-CoV-2. For tetramer staining, 1 million cells were transferred into 96-well V-bottom polystyrene plates and washed once with FACS buffer (PBS supplemented with 2% FBS and 2 mM EDTA), stained with 15 μg/ml the APC-labeled MHC class I tetramer for 1h at room temperature. Cells were then washed twice with FACS buffer, and stained for 30 min on ice with a cocktail of antibodies diluted in a 1:1 mixture of FACS buffer and the Brilliant Stain Buffer (BD Biosciences). The antibody cocktail consisted of APC/Cy7-CD3 (SK7, Biolegend), BD Horizon*ä* BV650-CD8 (RPA-T8, BD Biosciences), BD Horizon*ä* BUV737-CD4 (SK3, BD Biosciences), and the LIVE/DEAD Zombie Aqua*™* Fixable Viability Kit (Biolegend). After 3 additional washes with FACS buffer, cells were fixed and analyzed on an LSRFortessa_TM_ (BD Biosciences).

### Tetramerization of biotinylated MHC class I monomers with metal-labeled streptavidin

Streptavidin was labeled with metal as previously described (31). Biotinylated HLA-B*40:01 monomers with SARS-CoV-2 Nuc_322-331_ were obtained from the NIH Tetramer Core Facility. Tetramerization was performed as previously described (31). Briefly, the biotinylated monomers and metal-labeled streptavidin were each diluted to 50 μg/ml in PBS. A total of 10 μl of metal-labeled streptavidin was then transferred to 100 μl of peptide-MHC class I monomer. The solution was then mixed and incubated for 10 min at room temperature. After repeating the process twice (resulting in a total of 30 μl of metal-labeled streptavidin being transferred to 100 μl of the monomer solution), CyFACS buffer (metal contaminant-free PBS (Rockland) supplemented with 0.1% bovine serum albumin and 0.1% sodium azide) was added to reach a final volume of 500 μl. For each specimen containing up to 6x10_6_ cells, 100 μl metal-labeled tetramer was used.

### Preparation of specimens for CyTOF

Cryopreserved PBMCs were thawed and cultured overnight. Baseline specimens were stained directly in the absence of *ex vivo* stimulation. For identification of antigen-specific T cells through intracellular cytokine staining, 6x10_6_ cells were stimulated with 0.5 mg/ml anti-CD49d clone L25 (BD Biosciences) and 0.5 mg/ml anti-CD28 clone L293 (BD Biosciences) in the absence or presence of peptides for 4h in RP10 media (RPMI 1640 medium (Corning) supplemented with 10% fetal bovine serum (FBS, VWR), 1% penicillin (GIBCO), and 1% streptomycin (GIBCO)) in the presence of 3 mg/ml Brefeldin A Solution (eBioscience). Peptides used were 1 mM PepMix SARS-CoV-2 Peptide (Spike Glycoprotein) (JPT Peptide Technologies), 1 mM PepMix SARS-CoV-2 Peptide (NCAP) (JPT Peptide Technologies), or 1 mM PepMix SARS-CoV-2 Peptide (MEVTPSGTWL) (custom synthesized by JPT Peptide Technologies).

### CyTOF staining

We designed a 38-parameter CyTOF panel that allows assessment of the phenotypes, differentiation states, effector functions, and activation status of T cells, as well as homing receptors and transcription factors (Table S3). Antibodies that required in-house conjugation were conjugated to their corresponding metal isotopes using X8 antibody labeling kits according to manufacturer’s instructions (Fluidigm). For CyTOF staining, 6x10_6_ cells were blocked for 15 min on ice with sera from mouse (Thermo Fisher), rat (Thermo Fisher), and human (AB serum, Sigma-Aldrich) in Nunc_TM_ 96 Deep-Well polystyrene plates (Thermo Fisher). Cells were washed twice with CyFACS buffer, then stained with tetramer for 1h at room temperature in the presence of 50 nM dasatinib (Sprycel) to reduce T-cell receptor internalization and improve tetramer staining. Cells were then washed twice with CyFACS buffer, and stained for 45 min on ice with the cocktail of CyTOF surface staining antibodies (Table S3) in a total volume of 100 μl / well. Cells were then washed three times with CyFACS buffer and stained with Maleimide DOTA (Macrocyclics) for 30 min on ice. Cells were then washed twice with CyFACS and fixed overnight at 4°C with 2% PFA (Electron Microscopy Sciences) in metal contaminant-free PBS (Rockland). The next day, cells were permeabilized by incubation for 30 min at 4°C with fix/perm buffer (eBioscience), and then washed twice with Permeabilization Buffer (eBioscience). Cells were then blocked for 15 min on ice with sera from mouse (Thermo Fisher) and rat (Thermo Fisher). Cells were then washed twice with Permeabilization Buffer (eBioscience) and stained for 45 min on ice with the cocktail of CyTOF intracellular staining antibodies (Table S3). Cells were next washed once with CyFACS and incubated for 20 min at room temperature with 250 nM Cell-ID_TM_ DNA Intercalator-Ir (Fluidigm) in 2% PFA diluted in PBS. As a final step, cells were washed twice with CyFACS, once with Maxpar^®^ Cell Staining Buffer (Fluidigm), once with Maxpar^®^ PBS (Fluidigm), and once with Maxpar^®^ Cell Acquisition Solution (Fluidigm).

Immediately prior to acquisition, cells were resuspended to a concentration of 7 x10_5_ / ml in EQ_TM_ calibration beads (Fluidigm) diluted in Maxpar^®^ Cell Acquisition Solution. Cells were acquired at a rate of 250-350 events/sec on a CyTOF2 instrument (Fluidigm) at the UCSF Flow Cytometry Core.

### CyTOF data analysis

The CyTOF datasets were normalized to EQ_TM_ calibration beads using Fluidigm’s CyTOF software, exported as FCS files, and imported into FlowJo (BD) and Cytobank for gating and downstream analysis. Total T cells were identified using a sequential gating strategy based on DNA content, viability, cell length, and a CD3+CD19- gate. SARS-CoV-2-specific T cells were then identified by subgating on CD4+ or CD8+ T cells as appropriate, followed by tetramer+ or IFN*γ*+ gating. FlowJo (BD) was used for gating, generation of histogram plots, and mapping of defined populations onto the tSNE plots. Cytobank was used to calculate the median signal intensity (MSI) of cell populations based on standard two-dimensional dot plots. Cytobank was also used to generate tSNE plots and FlowSOM plots using the default settings (with a modification of total metaclusters from 10 to 5 for FlowSOM analysis). All of the phenotyping markers were used in tSNE and FlowSOM analysis, except for CD19, which was a parameter used in the upstream gating strategy. Line graphs were generated using ggplot2 in R. The raw CyTOF datasets generated from this study are available for download through the public repository Dryad via the following link: https://doi.org/10.7272/Q6D21VVD.

### Serology

The flow cytometry-based serological assay, based off previously validated methods (32), was used to quantitate the relative levels of IgA1, IgA2, IgE, IgG1, IgG2, IgG3, IgG4, and IgM against the nucleocapsid and various domains of the spike protein (S1, S2, and the RBD domain of S1). This assay for measuring serum antibody levels uses biotin-conjugated antigens coupled to streptavidin-coated microspheres (beads). Incubation of antigen-bead complexes with patient sera, and subsequent staining by fluorescently-conjugated, isotype-specific antibodies, produces a flow cytometric readout of bead fluorescence which reveals the levels of antigen-specific antibodies and their isotypes. The assay was calibrated using mouse monoclonal (IgG_2B_) antibodies raised against the RBD/S1/S2/NP antigens. The calibration revealed high specificity and no cross-reactivity between antigens, with the exception of cross-reactivity of anti-RBD antibodies against S1, which was expected as RBD is contained within S1. To assess isotype usage of RBD/S1/S2/NP-specific antibodies from PID4103, we incubated antigen-coated beads with plasma from the five timepoint specimens of PID4103 (1:2 dilution in HBSS + 0.1% BSA media). Non-specific antibody binding was assessed by incubation of plasma with uncoated (antigen-free) beads. The beads were washed with HBSS supplemented with 0.1% BSA, and then stained with an isotype-specific, fluorescently conjugated cocktail of antibodies at a concentration empirically determined to have minimal background binding to both antigen-coated and uncoated beads. Processed beads were analyzed using a BD FACSymphony_TM_ flow cytometer. Raw values were normalized by subtraction of non-specific signal as determined by the signal from the antigen-free beads. The grey shaded areas (Fig. 2G) highlight low-confidence signals where the mean fluorescence intensity (MFI) difference between specific and non-specific signal was less than 100 units.

### K-means cell clustering based on CyTOF profile

We performed unsupervised cell clustering of all the measured parameters using a k-means algorithm implemented using the kmeans function in R (https://www.R-project.org/). Each kind of measured parameter was centered using the mean of the corresponding levels across the five timepoints and scaled using the standard deviation of the values, before proceeding with unsupervised clustering. To find the optimal number of clusters, the gap statistic, a metric that evaluates clustering efficiency by comparing the sum of within-cluster distance from real data and null data (33), was implemented using the R library factoextra (https://CRAN.R-project.org/package=factoextra). The gap statistic was measured with K values ranging from 1 to 100. Since the gap statistic trend curve increased as K increased, the optimal K value was selected as that within 1 standard error from the first local maximum. The optimal K value was determined to be 5, and the validity of cell subtypes was examined by visualizing measured parameter patterns using a heatmap generated using pheatmap (http://CRAN.R-project.org/package=pheatmap).

## ACKNOWLEDGEMENTS

This work was supported by the Van Auken Private Foundation, David Henke, and Pamela and Edward Taft (N.R.R.); philanthropic funds donated to Gladstone Institutes by The Roddenberry Foundation and individual donors devoted to COVID-19 research (N.R.R.); the Program for Breakthrough Biomedical Research (N.R.R., E.G., S.L., J.V.), which is partly funded by the Sandler Foundation; Awards #2164 (N.R.R.), #2208 (N.R.R.), and #2160 (to S.L.) from Fast Grants, a part of Emergent Ventures at the Mercatus Center, George Mason University; NIH R01 AI123127-05S1 (E.G.); and Emory’s Lowance Center for Human Immunology (E.G.). We acknowledge the NIH DRC Center Grant P30 DK063720 and the S10 1S10OD018040-01 for use of the CyTOF instrument, and the NIH Tetramer Core Facility (contract number 75N93020D00005) for providing the SARS-CoV-2 tetramers and biotinylated monomers, and support from CFAR (P30AI027763), and the James B. Pendleton Charitable Trust. We thank Stanley Tamaki and Claudia Bispo for CyTOF assistance at the Parnassus Flow Core; Nandhini Raman and Jane Srivastava for assistance in flow cytometry at the Gladstone Flow Core; Jeff Milush and Norman Jones for assistance with the specimens at the Core Immunology Lab; Emory Pediatrics / Winship Flow Cytometry Core (access supported in part by Children’s Healthcare of Atlanta) and Emory Children’s Clinical and Translational Discovery Core for their support with flow cytometry-based serological experiments; Heather Hartig for help with recruitment; Min-Gyoung Shin and Reuben Thomas for help with the k-means clustering; Warner Greene for helpful feedback on the project; Françoise Chanut for editorial assistance; and Robin Givens for administrative assistance.

## AUTHOR CONTRIBUTIONS

T.M. designed and performed experiments, conducted data analyses, and helped put together the manuscript; H.R. established experimental protocols generated streptavidin reagents; B.B. performed experiments and conducted data analyses; M.M. processed and banked specimens, and generated scripts for data analysis; J.N., G.X., J.F., and A.G. processed and banked specimens; V.M. and G.G. conducted CHIRP participant interviews, enrollment, and specimen collection; E.G. designed protocols, conducted data analysis and performed supervision, and conceived ideas for the study; E.N. established protocols, provided reagents, helped with experimental design, and conceived ideas for the study; S.L. established the CHIRP cohort and conducted CHIRP participant interviews, enrollment, and specimen collection, and conceived ideas for the study; N.R.R. performed supervision, conducted data analyses, wrote the manuscript, and conceived ideas for the study,. All authors read and approved the manuscript.

## COMPETING FINANCIAL INTERESTS

The authors declare no competing financial interests.

## SUPPLEMENTAL FIGURE LEGENDS

**Figure S1.**
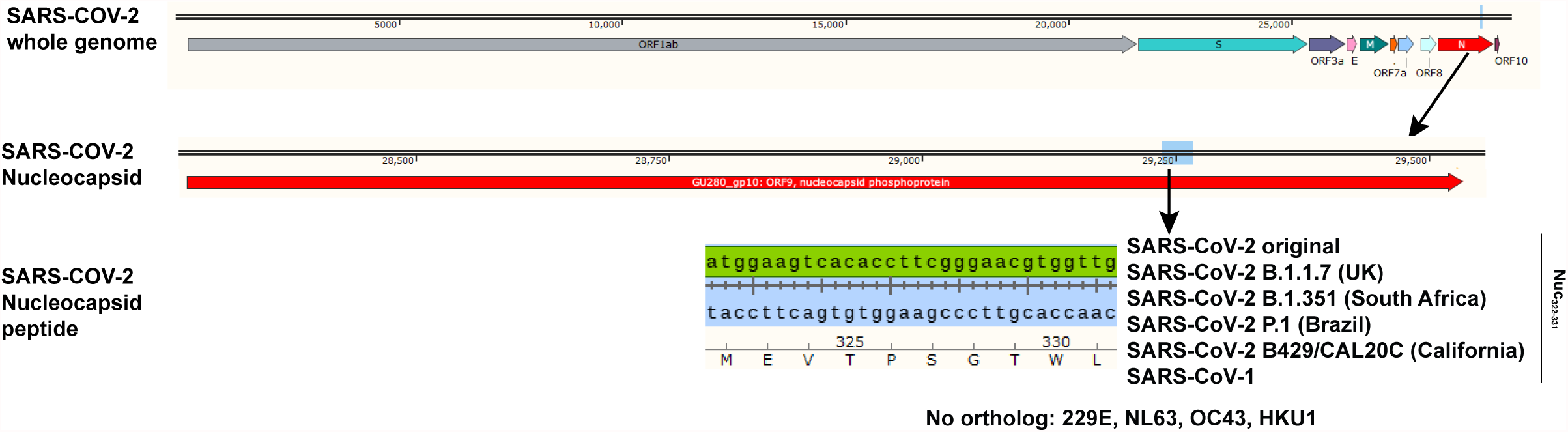
Nuc_322-331_ of SARS-CoV-2 is conserved in the SARS-CoV-2 variants-of-concern. Nuc_322-331_ resides near the C-terminus of the nucleocapsid protein of SARS-CoV-2, and its sequence, MEVTPSGTWL, is 100% conserved in the B.1.1.7, B.1.351, P.1, and B429/CAL20C first detected in the United Kingdom, South Africa, Brazil, and California, respectively. It is also 100% conserved in SARS-CoV-1, but no orthologs are present in the common cold coronavirus strains 229E, NL63, OC43, and HKU1. Shown are the location of the nucleocapsid gene within the SARS-CoV-2 genome, the location of the sequence encoding the peptide within the nucleocapsid gene, and both the nucleotide and amino acid sequences of corresponding to Nuc_322-331_.

**Figure S2.**
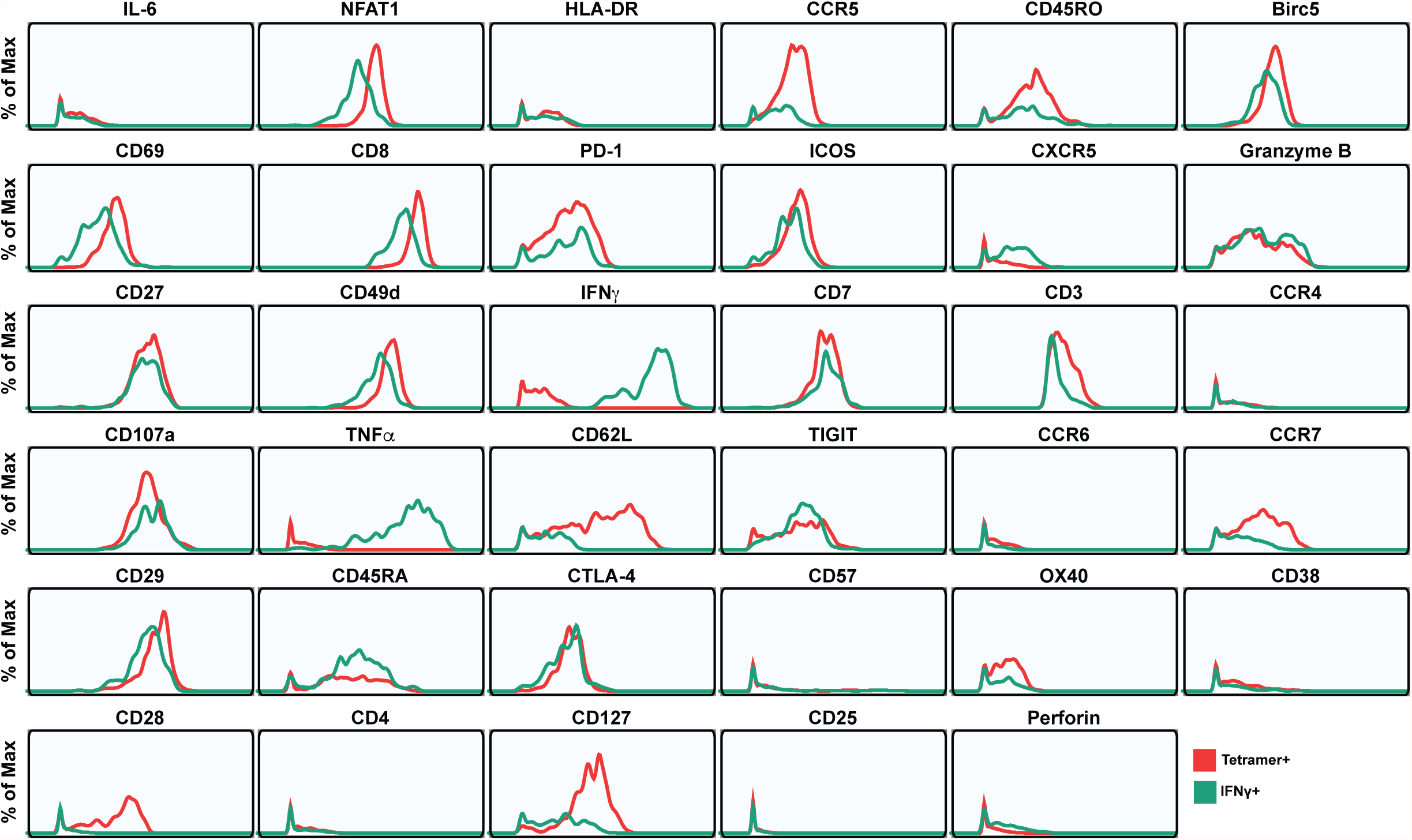
Expression levels of CyTOF antigens in Nuc_322-331_-specific CD8+ T cells from PID4103. The Nuc_322-331_-specific cells were identified either as unstimulated tetramer+ cells (*red*) or as cells producing IFN*γ* after peptide stimulation (*green*). For each antigen, the histograms represent merges of the five timepoints analyzed in this study.

**Figure S3.**
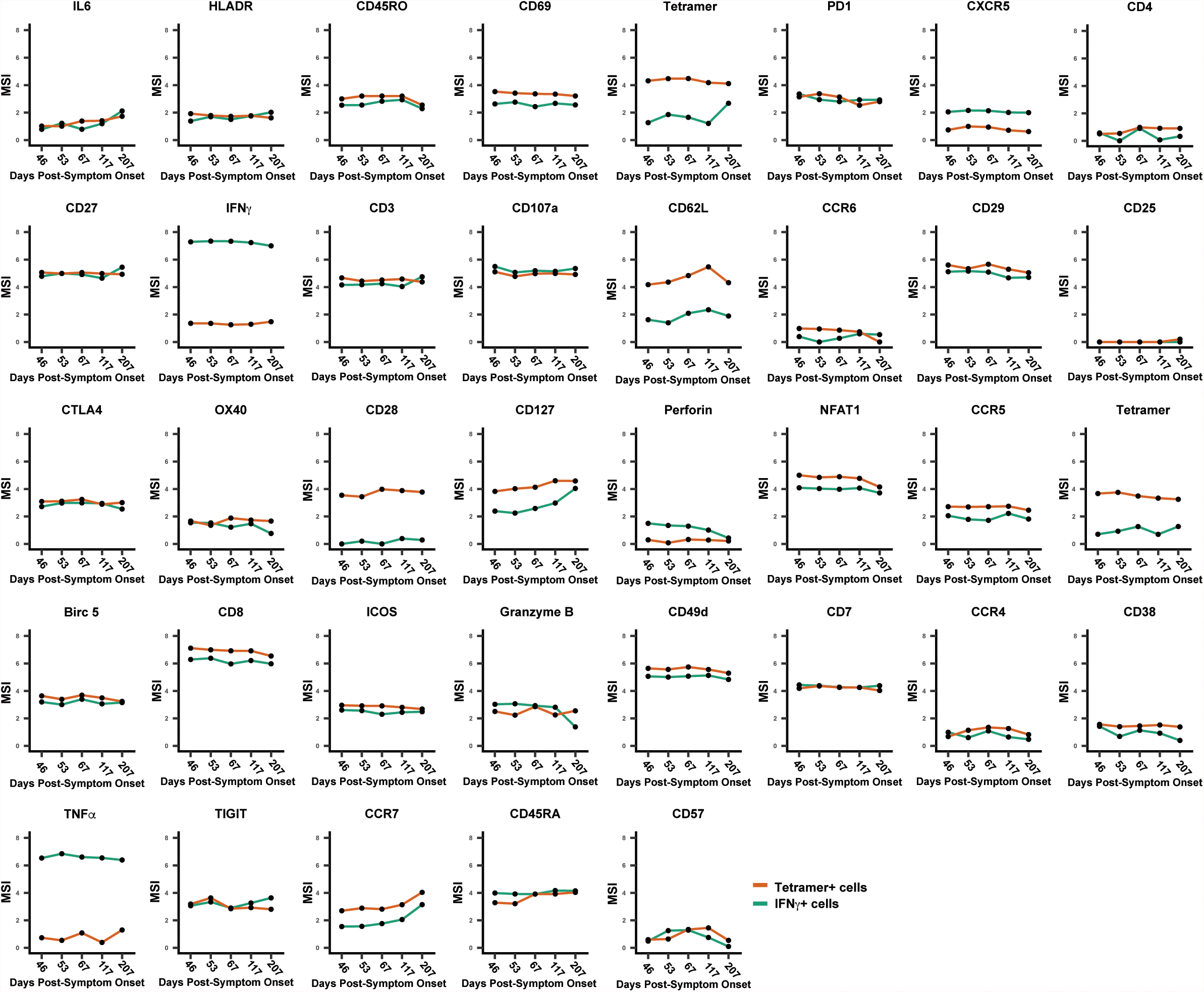
Expression levels of CyTOF antigens in Nuc_322-331_-specific CD8+ T cells from PID4103 as a function of time. Line graphs depict the antigens’ median staining intensities, as measured by CyTOF, among the tetramer+ cells in the unstimulated sample (*red*), and the IFN*γ*+ cells in the Nuc322-331- stimulated samples (*green*), at the five timepoints analyzed in this study.

**Figure S4.**
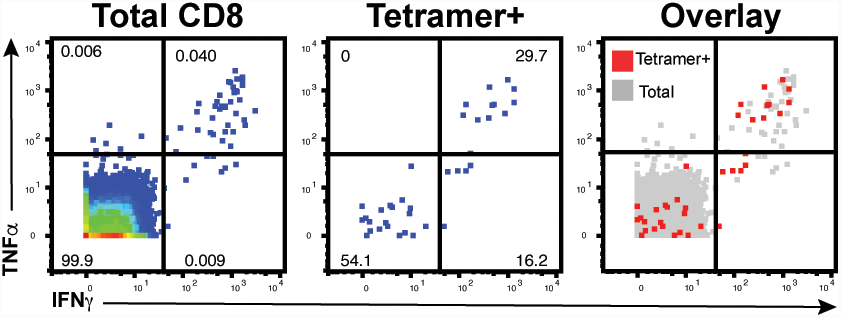
Approximately half of tetramer+ cells in Nuc_322-331_-stimulated samples do not secrete IFN*γ* or TNF*α*. PBMCs from PID4103 were stimulated with Nuc322-331, stained with HLA-B*40:01/Nuc322-331 tetramers, and analyzed by CyTOF. A total of 54.1% of tetramer+ cells expressed neither IFN*γ* nor TNF*α*, suggesting that approximately half of tetramer+ cells are not identified using the cytokine secretion assay.

**Figure S5.**
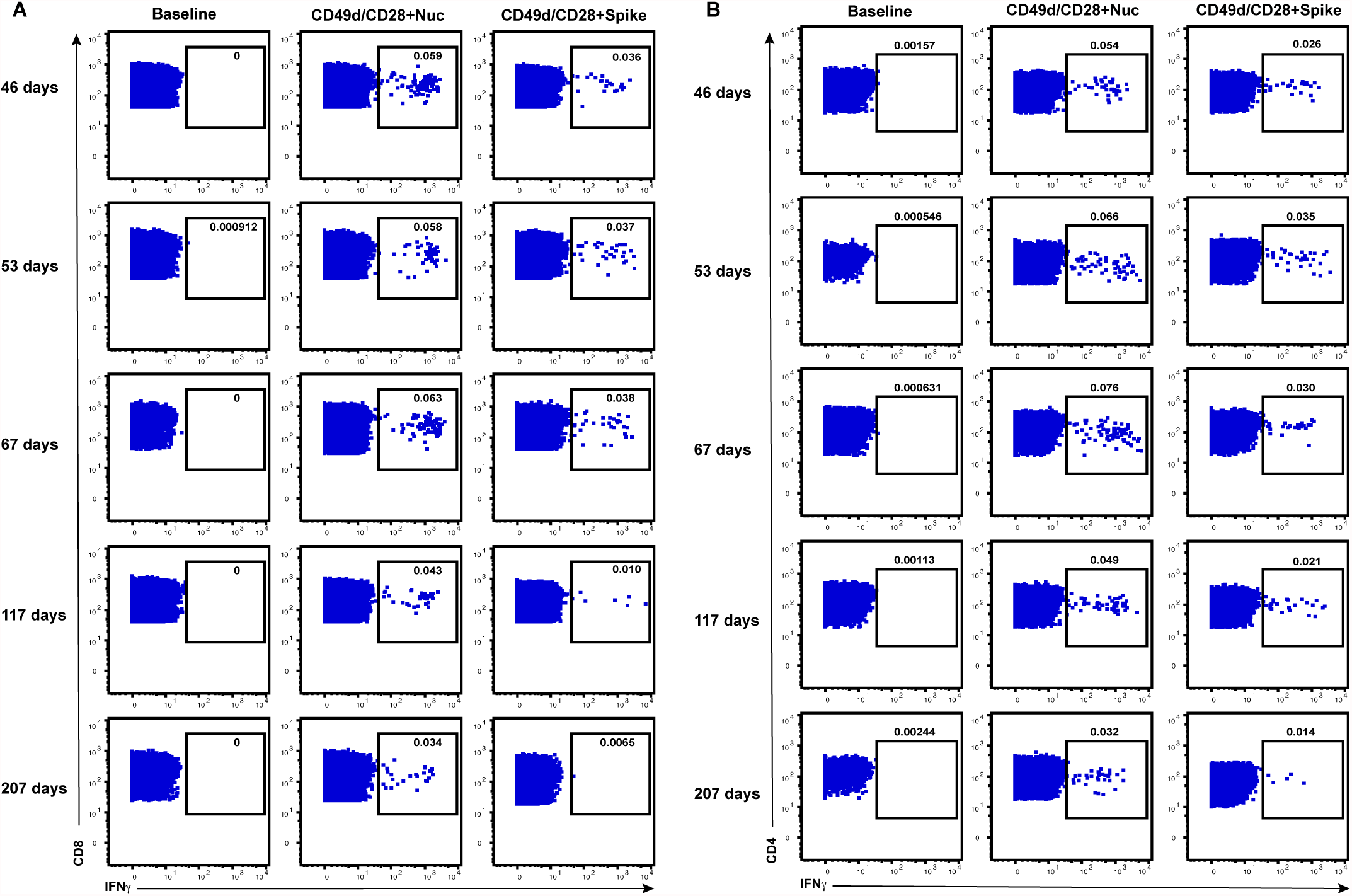
Longitudinal assessment of the CD8+ and CD4+ T cells of PID4103 directed against the nucleocapsid and spike proteins. CD8+ (A) and CD4+ (B) T cells specifically responding to stimulation with overlapping peptides spanning the entire nucleocapsid or spike proteins were identified by gating on the IFN*γ*+ cells. The cells were phenotyped by CyTOF at baseline, or following 4 hours of co-stimulation with *α*CD49d/CD28 and either the nucleocapsid or spike peptides. Stimulations were conducted in the presence of brefeldin A to enable detection of intracellular cytokines. Numbers correspond to the percentage of cells in each sample. Results are gated on live, singlet CD3+CD8+ cells (A) or live, singlet CD3+CD4+ cells (B). Timepoints correspond to days post symptom onset.

## SUPPLEMENTARY TABLES

**Table S1.**
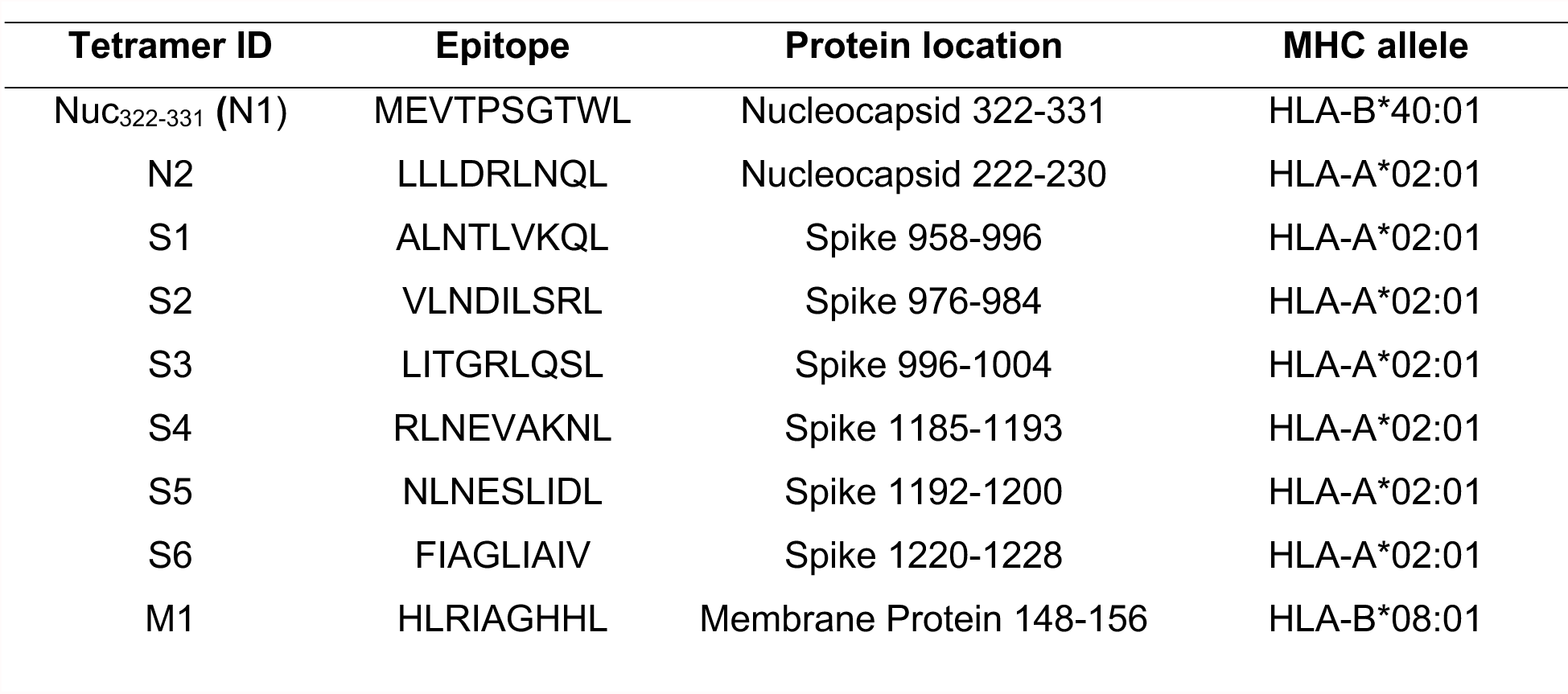
Tetramers screened by FACS.

**Table S2.** Participant Characteristics.

| **Patient ID** | **Gender** | **Age (years)** | **Date of symptom onset** | **Date of First PCR+ test** | **Date of Blood Draw(s)** | **Time Between Symptom Onset and analysis** | **Time Between first PCR+ test and analysis** |
| --- | --- | --- | --- | --- | --- | --- | --- |
| PID4103 | Female | 42 | 3/13/20 | 04/09/20 | 04/29/20<br>(PCR+) | 46 days | 20 days |
|  |  |  |  |  | 05/06/20<br>(PCR-) | 53 days | 27 days |
|  |  |  |  |  | 05/20/20<br>(PCR-) | 67 days | 41 days |
|  |  |  |  |  | 07/08/20<br>(PCR-) | 117 days | 90 days |
|  |  |  |  |  | 10/07/20<br>(PCR-) | 207 days | 181 days |

**Table S3.**
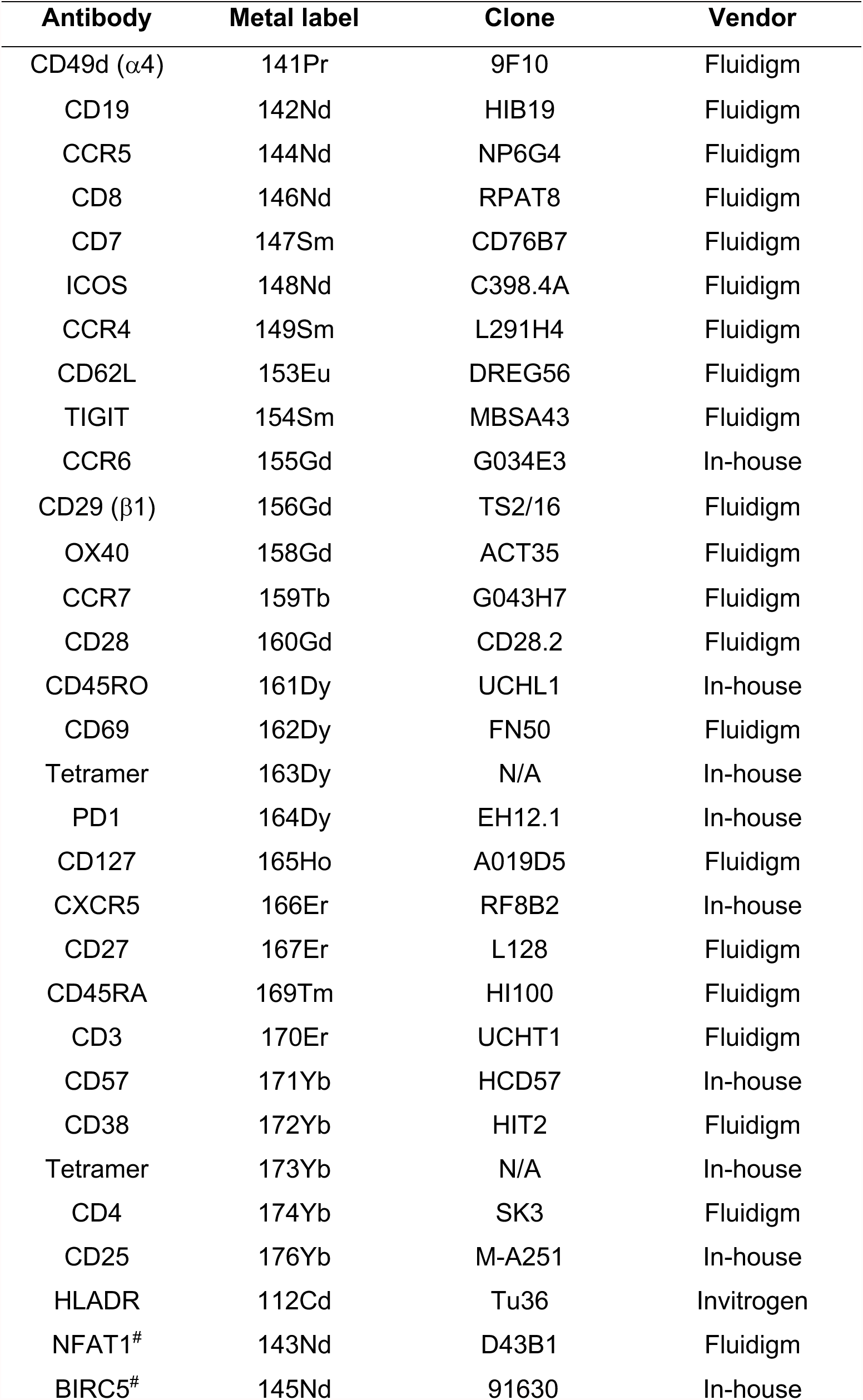
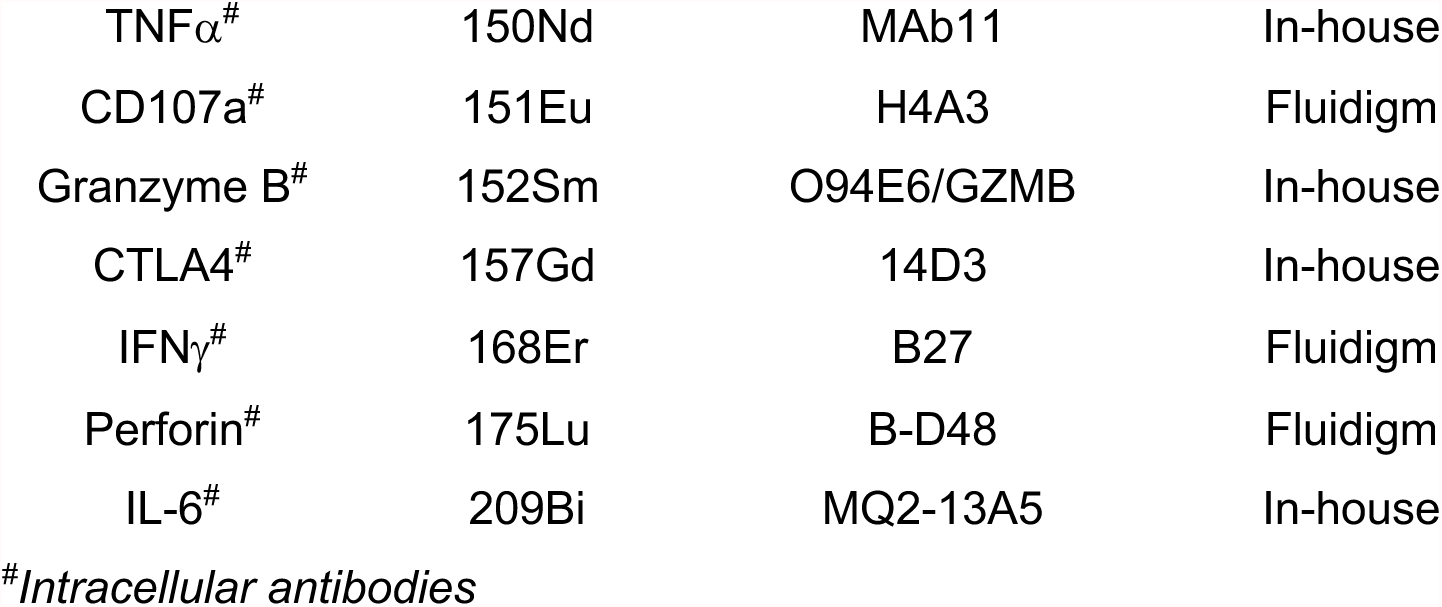
List of CyTOF staining antibodies.

## Notes

### Competing Interest Statement

The authors have declared no competing interest.

